# Ca^2+^-dependent phosphodiesterase 1 regulates the plasticity of striatal spiny projection neuron glutamatergic synapses

**DOI:** 10.1101/2024.04.24.590962

**Authors:** Shenyu Zhai, Shintaro Otsuka, Jian Xu, Vernon R. J. Clarke, Tatiana Tkatch, David Wokosin, Zhong Xie, Asami Tanimura, Hitesh K. Agarwal, Graham C. R. Ellis-Davies, Anis Contractor, D. James Surmeier

## Abstract

Long-term synaptic plasticity at glutamatergic synapses on striatal spiny projection neurons (SPNs) is central to learning goal-directed behaviors and habits. Although considerable attention has been paid to the mechanisms underlying synaptic strengthening and new learning, little scrutiny has been given to those involved in the attenuation of synaptic strength that attends suppression of a previously learned association. Our studies revealed a novel, non-Hebbian, long-term, postsynaptic depression of glutamatergic SPN synapses induced by interneuronal nitric oxide (NO) signaling (NO-LTD) that was preferentially engaged at quiescent synapses. This form of plasticity was gated by local Ca^2+^ influx through CaV1.3 Ca^2+^ channels and stimulation of phosphodiesterase 1 (PDE1), which degraded cyclic guanosine monophosphate (cGMP) and blunted NO signaling. Consistent with this model, mice harboring a gain-of-function mutation in the gene coding for the pore-forming subunit of CaV1.3 channels had elevated depolarization-induced dendritic Ca^2+^ entry and impaired NO-LTD. Extracellular uncaging of glutamate and intracellular uncaging of cGMP suggested that this Ca^2+^-dependent regulation of PDE1 activity allowed for local regulation of dendritic NO signaling. This inference was supported by simulation of SPN dendritic integration, which revealed that dendritic spikes engaged PDE1 in a branch-specific manner. In a mouse model of Parkinson’s disease (PD), NO-LTD was absent not because of a postsynaptic deficit in NO signaling machinery, but rather due to impaired interneuronal NO release. Re-balancing intrastriatal neuromodulatory signaling in the PD model restored NO release and NO-LTD. Taken together, these studies provide novel insights into the mechanisms governing NO-LTD in SPN and its role in psychomotor disorders, like PD.

## Introduction

The striatum promotes the selection of context-appropriate actions from a range of possibilities generated by the cortex.^1^ This selection process is shaped by reinforcement learning.^2–4^ Learning the association between context, action, and outcome manifests itself in the strength of the glutamatergic synapses on striatal spiny projection neurons (SPNs), the principal neurons of the striatum.^2,3,5–9^ Several forms of long-term synaptic plasticity have been reported in SPN synapses.^5,6,9,10^ But the relationship between them, their differential engagement, and how they might work together to shape learning remains a matter of speculation.

The best-studied form of striatal synaptic plasticity is endocannabinoid-mediated LTD (eCB-LTD). This form of plasticity is homosynaptic, postsynaptically induced, and presynaptically expressed.^11–15^ It is restricted to corticostriatal glutamatergic synapses and governed by G-protein coupled receptors (GPCRs) that are specific to direct pathway SPNs (dSPNs) and indirect pathway SPNs (iSPNs).^16–19^ In both types of SPN, eCB-LTD requires activation of CaV1 (L-type) voltage-gated Ca^2+^ channels (VGCCs), primarily those with a low-threshold, CaV1.3 pore-forming subunit.^14,20–22^

A less studied form of striatal LTD at SPN glutamatergic synapses is heterosynaptic and expressed postsynaptically. It is triggered by nitric oxide (NO) released by low-threshold spiking interneurons (LTSIs) that express neuronal nitric oxide synthase (nNOS).^18,23^ In SPNs, NO activates soluble guanylyl cyclase (sGC), which elevates cyclic guanosine monophosphate (cGMP); cGMP signaling triggers the endocytosis of synaptic α-amino-3-hydroxy-5-methyl-4-isoxazolepropionic acid (AMPA) receptors and LTD. In contrast to eCB-LTD, NO-LTD is inducible at both corticostriatal and thalamostriatal glutamatergic synapses in both types of SPN.^18^ However, fundamental questions remain about how this form of synaptic plasticity is regulated and how it contributes to remodeling of striatal circuits in health and disease states such as Parkinson’s disease (PD).

A regulated step in SPN NO signaling is the degradation of cGMP. This step is controlled by a group of enzymes called phosphodiesterases (PDEs). Two cGMP degrading PDEs – PDE1B and PDE10A – are highly enriched in the striatum.^24,25^ Although both PDEs can degrade cGMP and cyclic adenosine monophosphate (cAMP), their propensity to do so is not the same: PDE1B has a strong preference for cGMP, whereas PDE10A preferentially degrades cAMP.^26,27^ Unlike other PDEs, PDE1B is activated by Ca^2+^ and calmodulin,^28,29^ making it dependent upon synaptic activity and linking it to both eCB-LTD and long-term potentiation (LTP), which also rely upon an elevation in cytosolic Ca^2+^.^14,30^ However, the cellular determinants of PDE1B activation and its impact on NO-LTD have not been characterized in SPNs.

To help fill this gap, *ex vivo* mouse experiments were used to characterize the mechanisms controlling PDE1 activity and its impact on synaptic plasticity. These studies revealed that dendritic cGMP signaling in SPNs is potently regulated by PDE1B, whose activity is selectively stimulated by Ca^2+^ influx through CaV1.3 L-type channels. As these channels are prominent in the dendrites and spines of SPNs,^31^ this coupling restricted NO-LTD to quiescent dendrites and synapses – in sharp contrast to all other known forms of synaptic plasticity in SPNs. A gain-of-function mutation in the pore-forming subunit of the CaV1.3 channel led to enhanced dendritic Ca^2+^ entry in SPNs and disruption of NO-LTD. NO-LTD also was disrupted in a mouse model of PD, but in this case the deficit was attributable to a loss of LTSI NO generation; reversing the PD-induced neuromodulatory imbalance restored NO-LTD in SPNs. Taken together, these studies establish that PDE1 confers a novel, activity-dependence upon heterosynaptic NO-LTD in striatal SPNs, ensuring it is effectively coordinated with other forms of synaptic plasticity. Moreover, the PDE1 regulation of NO-LTD creates a mechanism by which global synaptic strength in SPNs can be maintained at the expense of inactive synapses, potentially contributing to behavioral flexibility.

## Results

### PDE1 regulated cGMP-dependent LTD in SPNs

Despite the robust expression of PDE1B in the striatum (Figure 1A),^24,25,32–35^ its functional role there is still unclear. Given that PDE1B has a high affinity for cGMP (relative to cAMP)^26,36,37^ and that cGMP-related signaling molecules are highly expressed in the striatum,^38–41^ one possible function of PDE1B is to regulate cGMP signaling in principal striatal SPNs. As a first step toward assessing this possibility, the expression of PDE1B in iSPNs and dSPNs was determined. Fixed slices from Drd1-tdTomato or Drd2-EGFP reporter mice^42^ were immunostained using a PDE1B-specific antibody. PDE1B immunoreactivity was evident in both iSPNs and dSPNs (Figures 1B and S1A).

**Figure 1.**
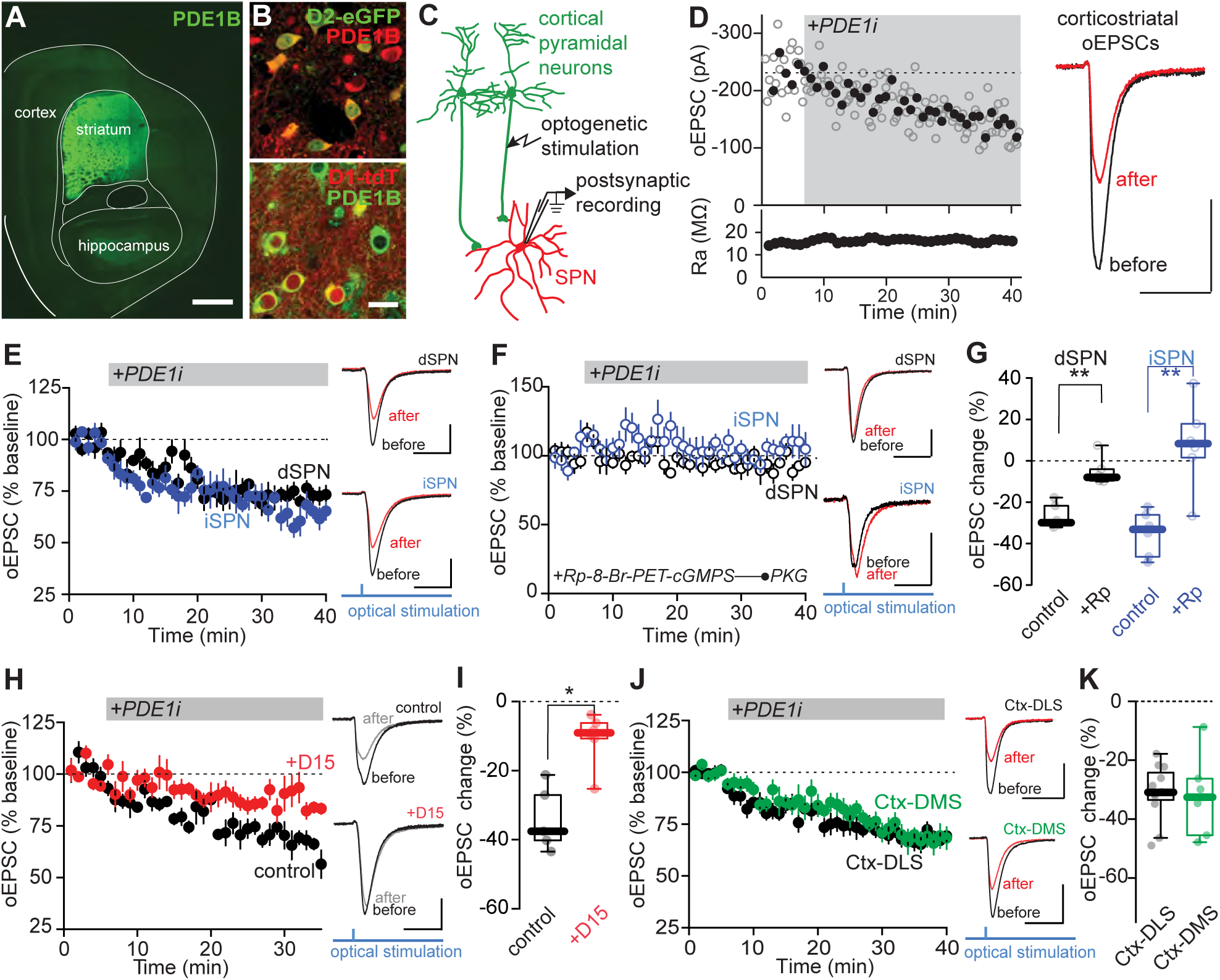
PDE1 inhibition leads to cGMP-dependent LTD at corticostriatal synapses in SPNs. (A) Composite confocal image showing strong expression of PDE1B in the striatum. Scale bar is 1 mm. (B) Confocal images showing colocalization of PDE1B immunoreactivity with D2-eGFP fluorescence (top) and D1-tdTomato (tdT) fluorescence (bottom). Scale bar is 20 μm. (C) Schematic diagram of the recording configuration. Whole-cell patch clamp recordings were made from SPNs in acute brain slices of Thy1-ChR2 mice and EPSCs were optogenetically evoked with brief blue LED pulses. (D) Sample whole-cell recording from an SPN upon bath application of specific PDE1 inhibitor AF64196 (PDE1i, 0.28 μM). Plots show amplitudes of optogenetically evoked EPSC (oEPSC) and access resistance (Ra) as a function of time. Filled symbols specify the averages of six data points. The dashed line represents the average EPSC amplitude before PDE1i application. The averaged EPSC traces before and 30 min after PDE1 inhibition are shown on the right. In this panel and panels (E), (F), (H) and (J), scale bars are 100 pA x 20 ms. (E) Left, application of PDE1i induced synaptic depression in the SPNs of the direct and indirect pathways. Time course of normalized EPSC in dSPNs (black, n = 6 neurons from 5 animals) and iSPNs (blue, n = 6 neurons from 5 animals) upon bath application of PDE1i is shown. Data are represented as mean ± SEM. Right, sample EPSC traces before and 30 min after PDE1 inhibition. (F) Left, intracellular dialysis of PKG inhibitor Rp-8-Br-PET-cGMPS (Rp, 10 mM) blocked PDE1i-induced LTD in both dSPNs (black, n = 6 neurons from 3 animals) and iSPNs (blue, n = 7 neurons from 4 animals). (G) Box plot summary of data in (E) and (F) showing changes in oEPSC (relative to baseline) from the last 5 min of recordings. **p < 0.01, Mann-Whitney test. (H) Left, endocytosis-disrupting peptide D15 (1 mM), when applied intracellularly, impairs PDE1i-LTD (n = 6). (I) Box plot summary of changes in oEPSC caused by PDE1i in the presence of intracellular D15 peptide and in interleaved controls. *p < 0.05, Mann-Whitney test. (J) Left, PDE1i induces similar synaptic depression of EPSC at cortex (Ctx)-dorsolateral striatum (DLS) synapses (n = 12 neurons from 10 animals) and Ctx-dorsomedial striatum (DMS) synapses (n = 7 neurons from 5 animals). Right, sample EPSC traces before and 30 min after PDE1 inhibition. (K) Box plot summary of data in (H) showing changes in oEPSC from the last 5 min of recordings.

Previous studies have implicated cGMP signaling in a postsynaptically expressed form of LTD at SPN glutamatergic synapses (NO-LTD).^18,23,43^ To determine if PDE1B regulated NO-LTD, whole-cell patch clamp recordings were made from SPNs in acute brain slices containing dorsolateral striatum (DLS) of Thy1-ChR2 mice, and corticostriatal excitatory postsynaptic currents (EPSCs) were optogenetically evoked with brief blue LED pulses (Figure 1C). Inhibition of PDE1 with a potent and specific inhibitor AF64196 (PDE1i, 0.28 µM)^44^ led to a significant time-locked depression of the evoked EPSC amplitude (Figure 1D). As expected from the distribution of PDE1B protein and the heterosynaptic nature of NO signaling, the synaptic depression induced by PDE1i was similar in iSPNs and dSPNs identified using the Drd1-tdTomato or Drd2-EGFP reporter lines (Figure 1E). Furthermore, the PDE1i-induced synaptic depression was persistent (or long-term), as it was sustained after washing out the inhibitor (Figure S1B). A second, less selective PDE1 inhibitor, 8-methoxymethyl-3-isobutyl-1-methylxanthine (8MM-IBMX, 10 μM), also induced a similar depression of corticostriatal EPSCs (Figure S1C).^23^ The synaptic depression induced by PDE1i was not dependent upon optical stimulation, as it could also be reproduced when EPSCs were evoked by electrical stimulation (Figure S1D).

To define the role of postsynaptic cGMP signaling in the PDE1i-induced effects, pharmacological tools were used. A principal target of cGMP signaling in SPNs is protein kinase G (PKG).^38^ Intracellular application of the competitive PKG inhibitor Rp-8Br-PET-cGMPS (10 μM) blocked PDE1i effects in both dSPNs and iSPNs (Figures 1F and 1G; dSPNs, p = 0.0022; iSPNs, p = 0.0047). Furthermore, as previously demonstrated for NO-LTD,^18^ disrupting AMPA receptor (AMPAR) endocytosis with intracellular infusion of D15 peptide (1 mM)^45^ blunted the effects of PDE1i (Figures 1H and 1I; p = 0.0087). Also, consistent with a postsynaptic locus of expression, PDE1i-induced LTD was not accompanied by changes in paired pulse ratio (Figure S1E), a hallmark of presynaptic eCB-LTD.^46^ Lastly, as with NO-LTD,^18^ PDE1i-induced LTD was unaffected by antagonism of cholinergic signaling through muscarinic or nicotinic receptors (Figure S1F).

To test the possibility that the PDE1i-induced depression of synaptic strength was mediated by allosteric stimulation of phosphodiesterase 2 (PDE2) and a reduction in cyclic adenosine monophosphate (cAMP) levels in SPNs,^47^ the impact of PDE1i on corticostriatal synaptic transmission was re-examined in the presence of PDE2 inhibitor Bay 60-7550 at a PDE2-selective concentration (40 nM).^48^ Contrary to previous work in cell culture,^47^ inhibition of PDE2 in *ex vivo* brain slices did not attenuate the PDE1i effects on optogenetically evoked EPSC in SPNs (Figure S1G).

To determine if there were regional differences in PDE1B regulation of synapses, the magnitudes and kinetics of PDE1i-induced LTD at corticostriatal synapses in dorsomedial striatum (DMS) SPNs were compared to those in DLS SPNs. The synaptic depression induced by PDE1i was similar in both striatal regions (Figures 1J and 1K, p = 0.6504).

Taken together, these data are consistent with the hypothesis that PDE1B constitutively suppresses NO-LTD at glutamatergic synapses on SPNs. To test the synapse specificity of the PDE1 regulation, glutamatergic neurons in the thalamic parafascicular nucleus (PFN) were induced to express channelrhodopsin-2 (ChR2) by injecting KH288-grp-Cre mice with a Cre-off ChR2-eYFP adeno-associated virus (AAV)^49^ (Figure 2A). Mice were sacrificed 4-5 weeks after viral injection, *ex vivo* brain slices prepared and SPNs patch clamped (Figure 2B). Application of PDE1i reliably induced depression of optogenetically evoked thalamostriatal EPSCs in DLS SPNs (Figures 2C and 2D), demonstrating that glutamatergic synapses arising from both the cortex and thalamus are regulated by PDE1B.

**Figure 2.**
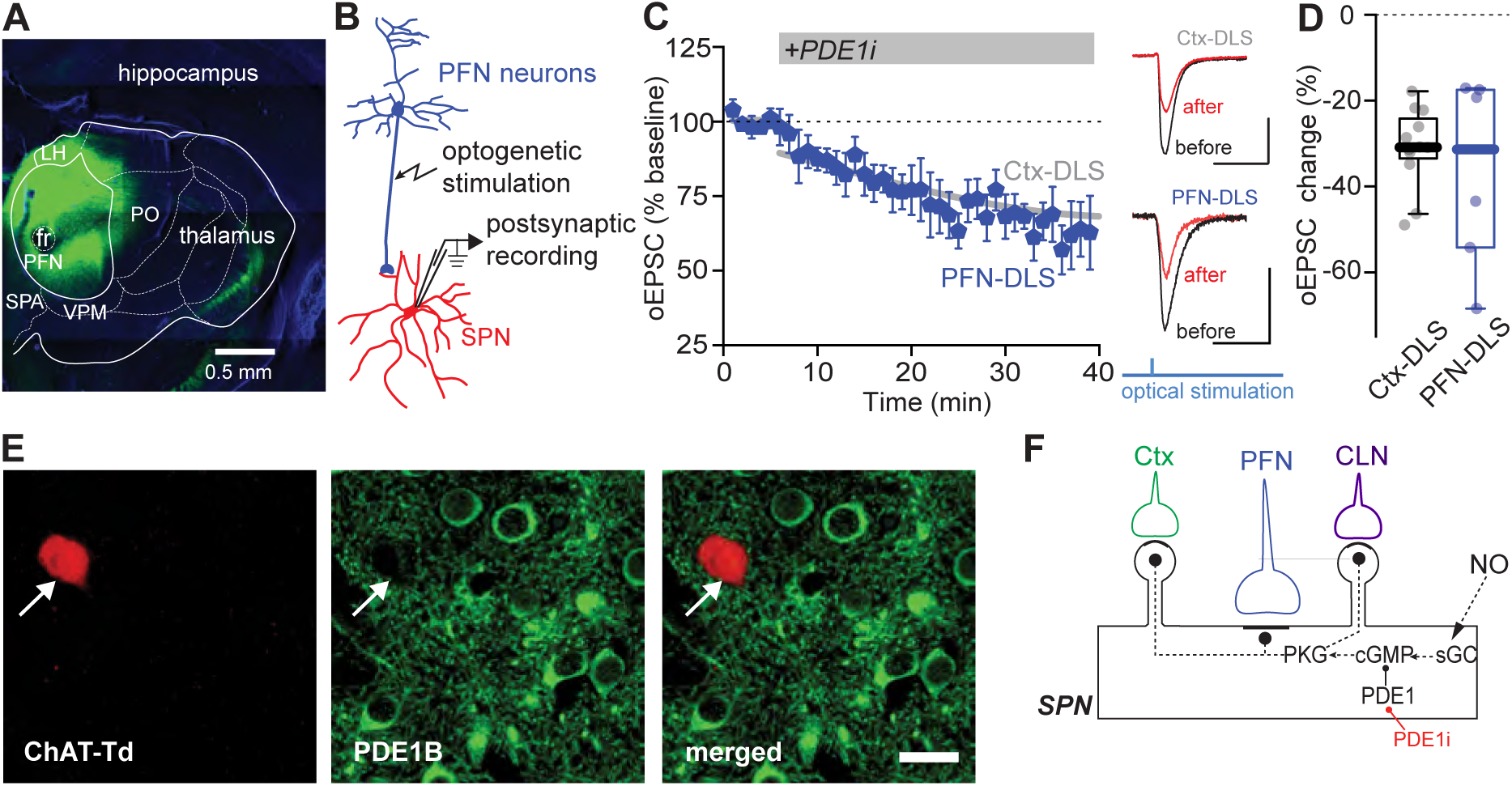
PDE1 inhibition leads to LTD at thalamostriatal synapses in SPNs. (A) Representative image of ChR2 expression in the parafascicular nucleus (PFN) of Grp-KH288 Cre mouse injected with Cre-off ChR2-eYFP. Scale bar is 0.5 mm. (B) Schematic diagram of the recording configuration. Whole-cell patch clamp recordings were made from SPNs in acute brain slices of Grp-KH288 Cre mice injected with Cre-off ChR2-eYFP. EPSCs were optogenetically evoked with brief blue LED pulses. (C) Left, PDE1i induces similar depression of EPSC at PFN-DLS synapses (n = 6 neurons from 4 animals) and Ctx-DLS synapses (fitted line, n=12). Right, sample EPSC traces before and 30 min after PDE1 inhibition. Scale bars are 100 pA x 20 ms. (D) Box plot summary of data in (C) showing changes in oEPSC from the last 5 min of recordings. (E) Representative confocal images showing lack of PDE1B immunoreactivity in ChIs labeled by tdTomato fluorescence (ChAT-tdT) in ChAT-Cre x Ai14 mice. Scale bar is 20 μm. (F) Schematic showing the modulation of NO-cGMP signaling in SPNs by PDE1i and downstream effect at corticostriatal and thalamostriatal synapses. CLN, centrolateral nucleus of the thalamus.

Striatal cholinergic interneurons (ChI) are also strongly excited by NO-induced cGMP signaling.^50–52^ However, it is not known whether cGMP in ChIs is regulated by PDE1B. In contrast to SPNs, PDE1B immunoreactivity was not detectable in ChIs identified by fluorescence in striatal sections taken from ChAT-Cre mice crossed with the Ai14 Cre-reporter line (Figure 2E). Moreover, inhibiting PDE1 had no effect on the spontaneous spiking activity of ChIs (Figure S2; p = 0.4779). Therefore, the primary function of PDE1 in the striatum is to suppress cGMP signaling and cGMP-dependent LTD at SPN glutamatergic synapses (Figure 2F).

In addition to inducing postsynaptic LTD, cGMP signaling can directly reduce dendritic CaV1 Ca^2+^ channel currents in SPNs.^18^ Indeed, bath application of the non-hydrolyzable cGMP analogue 8Br-cGMP (500 μM) diminished the dendritic Ca^2+^ transient evoked by a brief somatic depolarization from -60 mV to -40 mV (Figures S3A and S3B). To determine whether inhibition of PDE1 and elevation of in situ cGMP signaling would have a similar effect, PDE1i (AF64196, 0.28 μM) was bath applied and the experiment repeated. As predicted, application of PDE1i significantly diminished the depolarization-induced dendritic Ca^2+^ transient (Figures S3C and S3D), confirming that PDE1 is a crucial regulator of cGMP signaling in SPNs.

### PDE1 activity was dependent on CaV1.3 Ca^2+^ channels

Previous work in cell culture suggests that PDE1 is unique among phosphodiesterases in being activated by Ca^2+^-calmodulin.^36,53^ To determine the Ca^2+^-dependence of PDE1 activity in SPNs, neurons were loaded with the high affinity Ca^2+^ chelator 1,2-bis(o-aminophenoxy)ethane-N,N,N’,N’-tetraacetic acid (BAPTA, 5 mM) with the patch pipette. BAPTA dialysis significantly blunted the effects of PDE1i application (Figures 3A and 3C). This result suggests that Ca^2+^ chelation diminished PDE1B activity, occluding the effects of PDE1i. As previous work has implicated L-type CaV1 Ca^2+^ channels activated by synaptic depolarization in corticostriatal synaptic plasticity,^14,20^ a negative allosteric modulator of these channels (isradipine, 2 µM) was bath applied prior to PDE1i application. Doing so blunted the effects of the PDE1i on synaptic transmission (Figures 3B and 3C). Antagonizing N-methyl-D-aspartate (NMDA) receptors with D-AP5 (50 µM) or ryanodine receptors with dantrolene (10 µM) or calcium-permeable AMPARs (CP-AMPARs) with NASPM (20 μM) did not attenuate the PDE1i-mediated suppression of optogenetically evoked EPSCs in SPNs (Figures 3C and S4). Therefore, in SPNs, PDE1 is primarily activated by L-type Ca^2+^ channels.

**Figure 3.**
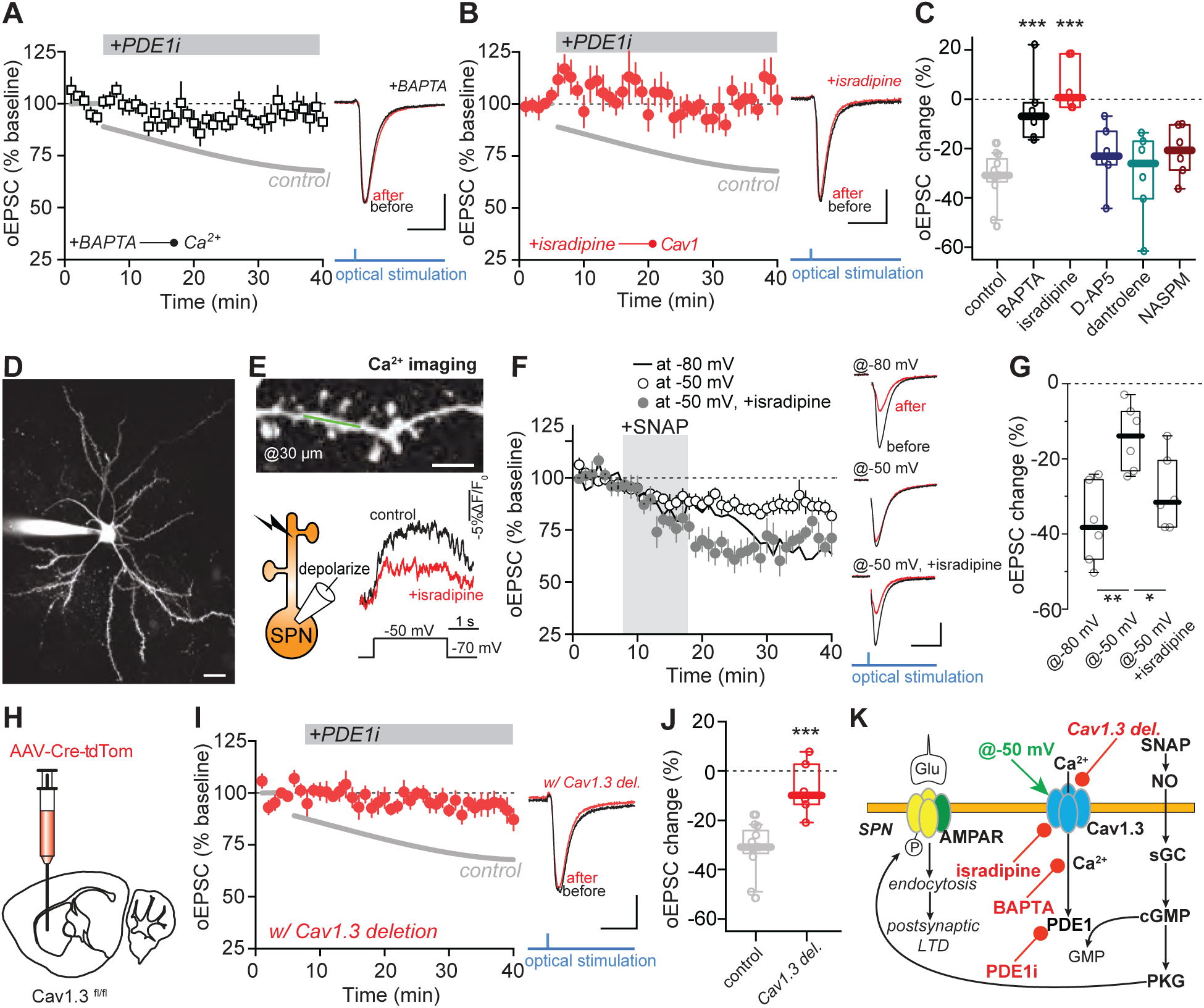
PDE1 activity depends on intracellular Ca^2+^ that comes from CaV1.3 Ca^2+^ channels. (A) PDE1i-LTD is blocked by intracellular BAPTA (5 mM, n = 6 neurons from 4 animals). Fitted historical control (n = 12 neurons from 10 animals) was shown as a grey line. Data are represented as mean ± SEM. Sample EPSC traces before PDE1i application and from the last 5 min of recording are shown on the right. In this panel and panels (B), (F) and (I), scale bars are 100 pA x 20 ms. (B) PDE1i-LTD is blocked by isradipine, a negative allosteric modulator of CaV1 calcium channels (2 μM, n = 6 neurons from 4 animals). (C) Box plot summary of changes in oEPSC (relative to baseline) induced by PDE1i application in the presences of various inhibitors. ***p < 0.001 compared to control, Mann-Whitney test. See also Figure S4. (D) Two-photon image of an SPN dialyzed with Fluo-4 and Alexa Fluor-568 through the patch pipette. Scale bar is 20 μm. (E) Top, two-photon image of a dendritic segment where line-scan Ca^2+^ imaging (indicated by a green line) was performed. Scale bar is 5 μm. Bottom left, schematic of dendritic Ca^2+^ imaging. Bottom right, representative Ca^2+^ signals evoked by depolarization from -70 mV to -50 mV before and after isradipine wash-in (1 μM) aligned with the voltage step protocol. See also Figure S5. (F) Time course of SNAP-LTD recorded with a holding potential of -80 mV, -50 mV or -50 mV in the presence of isradipine (-80 mV and -50 mV: both n = 6 neurons from 6 animals; -50 mV with isradipine: n = 6 neurons from 3 animals). Sample EPSC traces are shown on the right. (G) Box plot summary of data shown in (F). *p < 0.05, **p < 0.01, Mann-Whitney test. (H) Diagram showing injection of Cre recombinase-expressing AAV into the DLS of CaV1.3^fl/fl^ mice. (I) PDE1i-LTD is blocked by genetic knockdown of CaV1.3 (n = 6 neurons from 5 animals). (J) Box plot summary of data in (I) showing changes in oEPSC from the last 5 min of recordings. ***p < 0.001 compared to control, Mann-Whitney test. (K) Schematic showing the CaV1.3-PDE1 signaling pathway that negatively modulates postsynaptic LTD.

The dependence of PDE1 activity on depolarization-activated L-type Ca^2+^ channels led to the prediction that NO/cGMP-evoked LTD might be dependent on SPN activity state.^7,54^ *In vivo*, synaptic activity drives the membrane potential of SPNs from hyperpolarized ‘down-states’ to depolarized ‘up-states’ closer to the membrane potential where CaV1.3 channels open^31,55^ and, in principle, activate PDE1. In agreement with previous work showing dendritic localization of CaV1 channels in SPNs,^22,56^ a significant fraction of the dendritic Ca^2+^ signal evoked by stepping the somatic membrane potential from -70 mV to -50 mV (to mimic an up-state)^14,31^ was inhibited by bath application of isradipine (1 µM) (Figures 3D-3E and S5; p = 0.0004). To test the possibility that NO/cGMP-mediated synaptic plasticity would be attenuated during up-states by CaV1 channel-mediated activation of PDE1, the NO donor S-nitroso-N-acetylpenicillamine (SNAP) was used. When the somatic membrane potential was held at -80 mV (‘down-state’) to prevent CaV1 channel opening, SNAP application (100 µM, 10 min) induced a robust LTD at corticostriatal synapses (Figures 3F and 3G), as previously reported.^18,23^ However, when SPNs were held at -50 mV (‘up-state’), the LTD induced by SNAP was significantly reduced, but efficiently rescued by isradipine (Figures 3F and 3G; -80 mV vs. -50 mV, p = 0.0043; -50 mV vs. -50 mV + isradipine, p = 0.041), consistent with the hypothesis that Ca^2+^ influx through CaV1 channels activates PDE1 and suppresses NO-LTD.

Which subtype of CaV1 channel is linked to PDE1? As isradipine and other dihydropyridines do not distinguish channels with a CaV1.2 or CaV1.3 pore-forming subunit,^55^ the specific role of CaV1.3 channels was determined by genetic methods. In mice with the *Cacna1d* gene (which codes for the pore-forming subunit of CaV1.3 channels) floxed (see Experimental Procedures), an AAV expressing Cre recombinase and tdTomato was injected into the striatum (Figure 3H); four weeks later, *ex vivo* brain slices were prepared and tdTomato expression used to guide patch clamp recording. In Cre recombinase-expressing SPNs in which *Cacna1d* had been deleted (Figure S6), PDE1i application had no effect on corticostriatal synaptic transmission (Figures 3I and 3J). Taken together, these results suggest that Ca^2+^ flux through CaV1.3 channels stimulates PDE1, blunting cGMP-dependent postsynaptic LTD (Figure 3K).

### The gain-of-function G407R mutation in CaV1.3 channels impaired NO-LTD in SPNs

The involvement of CaV1.3 Ca^2+^ channels in the regulation of cGMP signaling has implications for our understanding of disease states with striatal determinants. Several missense mutations in the *Cacna1d* gene have been discovered in patients with neurodevelopmental disorders.^57^ One of these that has been associated with autism spectrum disorder (ASD) is a *de novo* glycine to arginine mutation at site 407 (G407R).^58^ This mutation has been reported to slow channel inactivation in cultured cells, increasing Ca^2+^ flux.^58^ To study the cellular consequences of this mutation, a knock-in mouse with the G407R mutation in the *Cacna1d* gene was created (Figures S7A-S7C; also see Experimental Procedures) and crossed into the D1-tdTomato reporter line to allow identification of dSPNs. Whole-cell voltage-clamp recording from identified dSPNs was performed in G407R heterozygous (*Cacna1d*^G407R/+^) and wildtype littermate mice in conjunction with two-photon Ca^2+^ imaging of the dendrites (Figure 4A). Ca^2+^ transients, indicated by increases in the fluorescence of Fluo-4, were evoked by depolarizing the soma of SPNs from -60 mV to -40 mV for 1 s. The contribution of CaV1 channels was assessed by bath application of isradipine (2 µM). As predicted from previous work, the dendritic Ca^2+^ transient attributable to Cav1 channels was significantly larger in *Cacna1d*^G407R/+^ dSPNs (Figures 4B and 4C; p = 0.0011), demonstrating that the G407R mutation increases Ca^2+^ entry through dendritic CaV1.3 channels.

**Figure 4.**
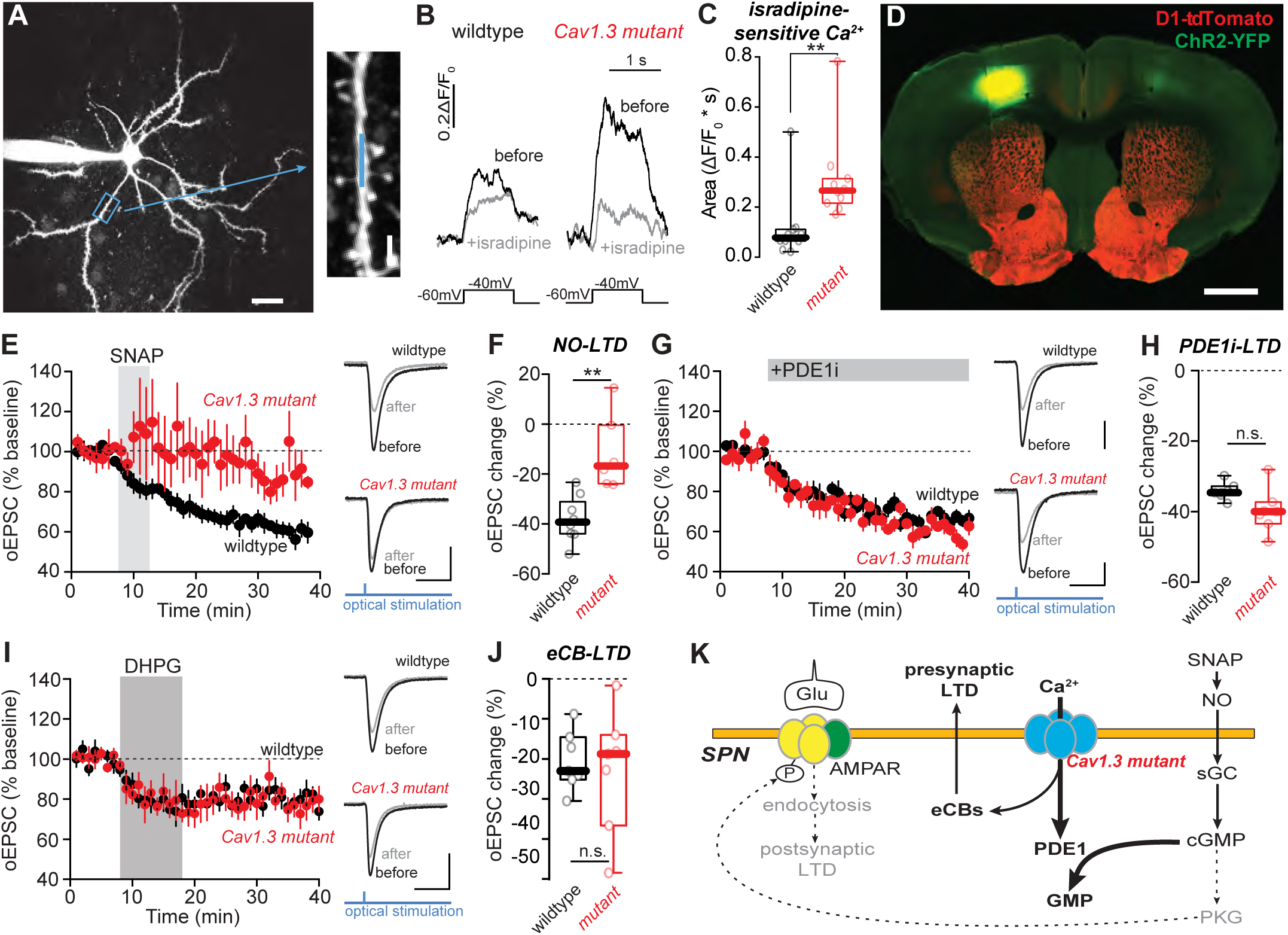
In mice with an ASD-associated G407R mutation in CaV1.3 channel, abnormal Ca^2+^ signal leads to impaired NO-LTD. (A) Schematic depicting the Ca^2+^ imaging assay. Left, a dSPN was patched in the whole-cell configuration, dialyzed with Ca^2+^-insensitive Alexa Fluor-568 and Ca^2+^-sensitive Fluo-4. Dendritic Ca^2+^ transients were triggered by somatic depolarization (from -60 mV to -40 mV for 1 s) through the patch pipette. Two-photon line-scan imaging was performed at segments of dendrites ∼45 µm from the soma (indicated by the green box). Right, a high-magnification image of a segment of dendrite. Line scan was indicated by the green line. Scale bars, 20 μm (left) and 2 μm (right). (B) Examples of dendritic Ca^2+^ transients before and after application of isradipine (2 μM) in wildtype siblings or *Cacna1d*^G407R/+^ mice were temporally aligned with the voltage step protocol. (C) Box plot summary of isradipine-sensitive component of dendritic Ca^2+^ transient in wildtype and *Cacna1d*^G407R/+^ mice (both n = 10 cells from 4 mice). G407R mutation significantly increases depolarization-evoked Ca^2+^ transients in the dendrites. **p = 0.0011, Mann-Whitney test. (D) Representative confocal image of ChR2 expression in the motor cortex of D1-tdTomato mouse. Scale bar is 1 mm. (E) SNAP-LTD is impaired in *Cacna1d*^G407R/+^ mice. EPSC was evoked by wide-field blue LED illumination (0.3 ms). LTD was induced by bath application of SNAP (5 μM) for 5 min (indicated by the grey vertical bar). Plot shows EPSC amplitude as a function of time. Data are mean ± SEM. n = 8 dSPNs from 6 wildtype mice and n = 6 dSPNs from 4 *Cacna1d*^G407R/+^ mice. Scale bars in (E), (G) and (I) are 200 pA x 20 ms. (F) Box plot summary of LTD amplitudes from the last 10 min of recordings shown in (E). In *Cacna1d*^G407R/+^ mice, SNAP-induced LTD was significantly impaired. **p < 0.01, Mann-Whitney test. (G) PDE1i-LTD is normal in *Cacna1d*^G407R/+^ mice. Plot shows EPSC amplitudes as a function of time. Data are mean ± SEM. Both n = 7 neurons from 4 animals. (H) Box plot summary of PDE1i-LTD recordings shown in (G). PDE1i induced similar synaptic depression at corticostriatal glutamatergic synapses in *Cacna1d*^G407R/+^ mice and interleaved wildtype littermates. n.s. not statistically significant, Mann-Whitney test. (I) DHPG-LTD is similar in wildtype and *Cacna1d*^G407R/+^ mice. Here, DHPG-LTD was induced by DHPG (50 μM) application for 10 min (indicated by the grey vertical bar) at a holding potential of -60 mV in the presence of 1.2 mM Ca^2+^ in the bath solution. Data are mean ± SEM. Both n = 7 dSPNs from 5 mice. See also Figures S8. (J) Box plot summary of LTD amplitudes from the last 10 min of recordings shown in (I). The near-threshold DHPG-LTD protocol induced similar synaptic depression in wildtype and *Cacna1d*^G407R/+^ mice. n.s. not statistically significant, Mann-Whitney test. (K) Schematic showing that abnormal activity of the CaV1.3-PDE1 signaling pathway in the SPNs of *Cacna1d*^G407R/+^ mice impairs postsynaptic LTD.

Next, the functional implications of the mutation were determined by assessing the ability of NO to induce LTD in slices from *Cacna1d*^G407R/+^ mice. In principle, elevated Ca^2+^ entry through CaV1.3 channels in *Cacna1d*^G407R/+^ SPNs should impair NO-induced LTD by activating PDE1B. To restrict the analysis to corticostriatal synapses, an AAV carrying a ChR2 expression construct was injected into the motor cortex of *Cacna1d*^G407R/+^ mice and wildtype littermates approximately 3 weeks prior to preparing *ex vivo* brain slices (Figure 4D); optogenetic methods were then used to evoke EPSCs in dSPNs. To induce NO-LTD, the NO donor SNAP (5 μM) was applied for 5 min while SPNs were voltage-clamped at -70 mV. This protocol yielded robust LTD of optogenetically evoked EPSCs in dSPNs of wildtype siblings (62 ± 1% of baseline at 30–40 min), but only a weak depression of EPSCs in *Cacna1d*^G407R/+^ dSPNs (84 ± 4% of baseline at 30–40 min; Figures 4E and 4F; p = 0.0027, Mann-Whitney U test). In contrast, bath application of a PDE1 inhibitor induced a robust LTD in *Cacna1d*^G407R/+^ SPNs (60 ± 2% of baseline at 30–40 min) just as in wildtype littermates (66 ± 1% of baseline at 30–40 min) (Figures 4G and 4H). This suggests that the impairment of NO-LTD in *Cacna1d*^G407R/+^ SPNs was not caused by any changes in the postsynaptic machinery underlying LTD or LTD saturation by ambient NO, but rather was attributable to abnormally high PDE1 activity (Figure 4K). In agreement with this conclusion, striatal PDE1B protein levels were similar in wildtypes and mutants whether assessed by immunohistochemistry (Figure S7D) or Western blot (Figure S7E), arguing that the impairment in NO-LTD was not a consequence of a secondary alteration in SPN PDE1B expression.

CaV1.3 channels also have been implicated in postsynaptically induced, but presynaptically expressed, eCB-LTD at corticostriatal synapses in SPNs.^14,15,20,59,60^ To determine whether eCB-LTD was altered in *Cacna1d*^G407R/+^ dSPNs, the Group I mGluR agonist (S)-3,5-dihydroxyphenylglycine (DHPG; 50 μM) was bath applied for 10 minutes while holding dSPNs at a modestly depolarized potential (-50 mV), a manipulation that elicits eCB release from SPNs and triggers eCB-LTD.^14,15,19^ This protocol led to the induction of a robust LTD in dSPNs in both wildtype and *Cacna1d*^G407R/+^ mice (wildtype, 61 ± 0.7% of baseline at 31-40 min; *Cacna1d*^G407R/+^, 64 ± 1.3% of baseline at 31-40 min; Figure S8A). This was also the case when eCB-LTD was induced by recording at a less depolarized holding potential (-60 mV) (Figure S8B). To determine if there was a ‘ceiling effect’ on eCB-LTD, the extracellular Ca^2+^ concentration was lowered to 1.2 mM (from 2 mM). The magnitude of eCB-LTD was reduced by lowering extracellular Ca^2+^ concentration, but to a similar extent in both wildtype and *Cacna1d*^G407R/+^ mice (Figures 4I and 4J; p = 0.96). These results suggest that while Ca^2+^ entry through CaV1.3 channels is necessary to trigger eCB-LTD, the magnitude of the plasticity is controlled by other factors.

### Dendritic cGMP-evoked LTD was negatively regulated by cytosolic Ca^2+^

Our results suggest that peri-synaptic Ca^2+^ entry through CaV1.3 channels activates PDE1, suppresses cGMP signaling, and thus prevents the induction of postsynaptic LTD in SPNs. To test this hypothesis, a novel, cell-impermeable caged cGMP compound, DEAC450-cGMP,^52^ was used to control the spatiotemporal profile of cGMP signaling in SPNs. A patch pipette was used to dialyze SPNs with DEAC450-cGMP (75 μM), and a high concentration of the Ca^2+^ chelator ethylene glycol-bis(β-aminoethyl ether)-N,N,Nʹ,Nʹ-tetraacetic acid (EGTA, 5 mM) was used to blunt activation of PDE1 by CaV1.3 channel opening during synaptic stimulation produced by electrical stimulation of corticostriatal axons (Figures 5A and 5B). Intracellular uncaging of cGMP triggered a robust postsynaptic LTD (Figure 5C). This LTD was occluded by preincubation with SNAP (Figure S9A), suggesting that this DEAC450-cGMP-uncaging induced a form of LTD that relied upon the same signaling pathway as NO-LTD. In agreement with a postsynaptic locus of the LTD, the change in EPSC amplitude induced by DEAC450-cGMP uncaging was not accompanied by a change in PPR (Figure 5D). To assess the Ca^2+^ dependence of the LTD induced by intracellular cGMP uncaging, SPNs were dialyzed with a low concentration of EGTA (0.25 mM) enabling Ca^2+^ entry through CaV1.3 channels to activate PDE1. As expected, enabling PDE1 activation prevented photolysis of DEAC450-cGMP from inducing LTD (Figure 5C). Dialysis with DEAC450-cGMP and EGTA (0.25 mM) alone (i.e. without blue light exposure) had no effect on corticostriatal EPSC amplitude (Figure S9B).

**Figure 5.**
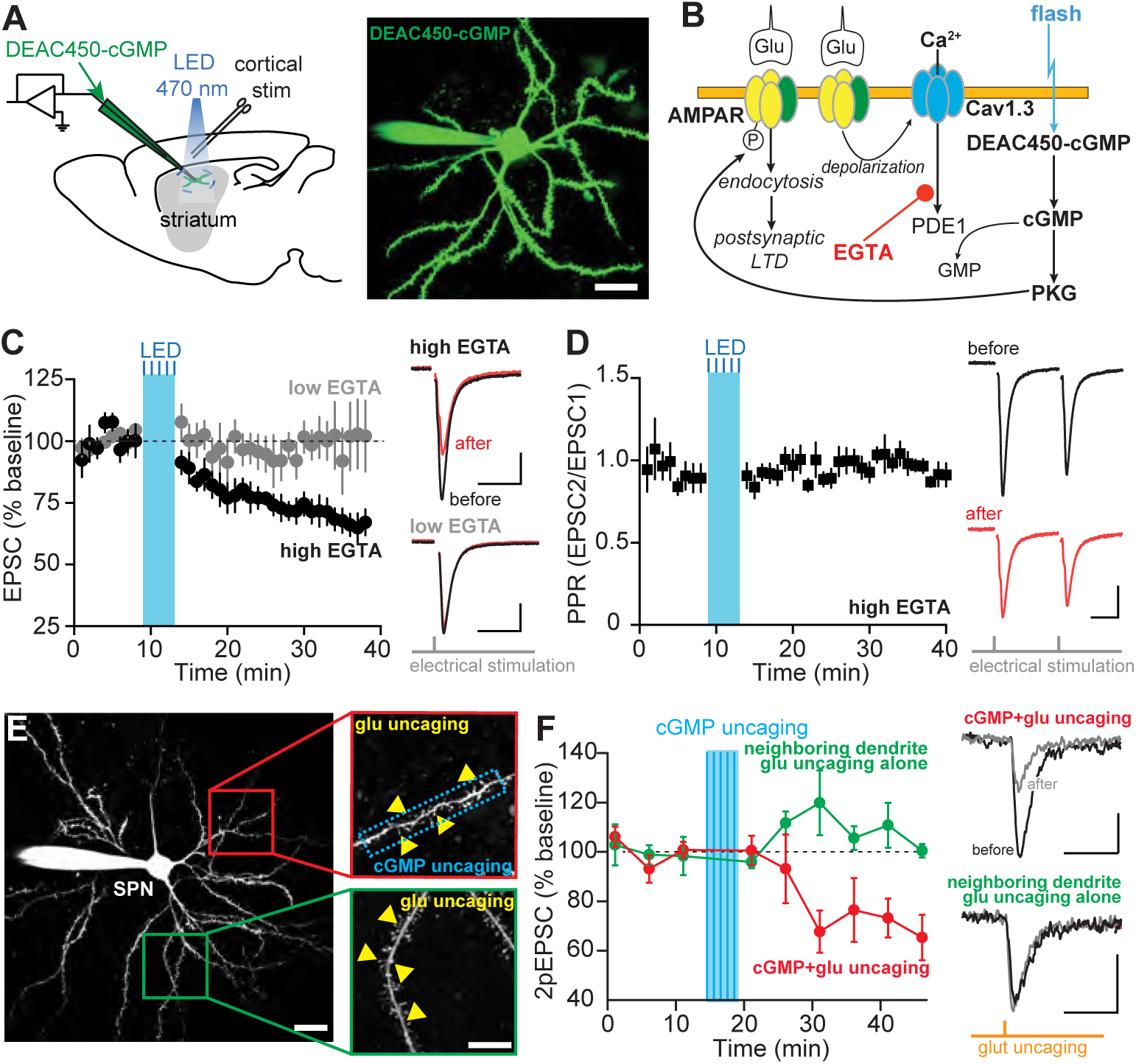
NO/cGMP-induced LTD is negatively regulated by intracellular Ca^2+^. (A) Left, schematic of experimental configuration for intracellular cGMP uncaging. Right, two-photon image of an SPN filled with caged cGMP (DEAC450-cGMP) through the patch pipette. Scale bar is 20 μm. (B) Schematic showing that induction of postsynaptic LTD by cGMP uncaging is only permitted when intracellular Ca^2+^ is dampened by high EGTA concentration. (C) Time course of electrically evoked EPSC before and after wide-field illumination with 470 nm LED (200ms every min, repeated five times) in SPNs intracellularly filled with DEAC450-cGMP (75 μM) supplemented with 5 mM EGTA (“high EGTA”, n = 7 neurons from 6 mice) or 0.25 mM EGTA (“low EGTA”, n = 6 neurons from 5 mice). In panels (C) and (D), scale bars are 100 pA x 20 ms. (D) Time course of paired-pulse ratio (PPR) of EPSC in the SPNs with high EGTA (the same cells as in (C)). (E) Experimental design of dual-color uncaging. Two-photon fluorescence images of an SPN (left) and its dendritic segments (right) are shown. SPNs were filled with 50 μM Alexa Fluor-594 and 75 μM DEAC450-cGMP supplemented with 5 mM EGTA. The synaptic strengths of dendritic spines were assayed by glutamate (glu) uncaging with 1-ms pulses of 720 nm two-photon laser (yellow triangles). cGMP uncaging was performed either on the same dendrite (upper right) with a 473 nm laser that covered a ∼35 μm dendritic segment (blue rectangle) or on a different dendrite (lower right). Scale bars are 20 μm (left) and 10 μm (right). (F) Left, plot of normalized two-photon glutamate uncaging-evoked EPSCs (2pEPSCs) as a function of time with localized cGMP uncaging on the same dendrite (n = 5 neurons from 3 animals) or a different dendrite (n = 4 neurons from 3 animals). Right, representative traces of 2pEPSCs before (black) and 20 min after cGMP uncaging (grey) in the same dendrite or in a different dendrite. Scale bars are 10 pA x 50 ms.

Dendrites can serve as independent signal processing units, performing local electrical and biochemical integration.^61–63^ To determine if cGMP signaling was localized, DEAC450-cGMP uncaging was restricted to a single dendritic segment, and the effects on synaptic strength were assessed using two-photon glutamate uncaging at that site and on a neighboring dendrite. This was possible because the absorption spectra of DEAC450-cGMP and MNI-glutamate do not overlap.^52,64,65^ To test the spatial extent of cGMP signaling, SPNs were first filled with DEAC450-cGMP and Alexa Fluo-594 through the patch pipette. Dendritic branches and spines were visualized with two-photon imaging by exciting Alexa Fluor-594 with 1040 nm light, a wavelength that does not photolyze DEAC450-cGMP or MNI-glutamate. MNI-glutamate was then uncaged with brief pulses of 720 nm two-photon light targeted at co-planar dendritic spine heads (Figure 5E). After obtaining baseline two-photon uncaging-evoked EPSCs (2pEPSCs), DEAC450-cGMP was uncaged in a short stretch (∼35 μm) of dendrite using 473 nm laser (200-ms pulses, repeated five times with 1-min intervals). Along the dendrite where cGMP had been uncaged, axospinous synapses underwent LTD (Figure 5F), as expected from the results above. However, on a neighboring dendrite, there was no change in 2pEPSC amplitude (Figure 5F). Dialysis with DEAC450-cGMP alone did not affect the 2pEPSC amplitude, nor did 473 nm laser stimulation of dendrites in the absence of DEAC450-cGMP (Figure S9C). Thus, intracellular cGMP signaling, which is tightly regulated by PDE1 activity, produced a dendritically localized, postsynaptic LTD.

A closely related question is how the CaV1.3 channel dependence of PDE1 limits cGMP signaling when dendrites are more strongly activated and competing forms of plasticity might be induced. Spatiotemporally convergent glutamatergic input to distal SPN dendrites can generate an NMDAR-dependent spike, creating the conditions necessary for LTP induction.^7,66,67^At the dendritic site of spike initiation, the depolarization is presumed to be strong enough to activate voltage-dependent Ca^2+^ channels, including low-threshold CaV1.3 channels,^66,68^ but how the spike-initiated depolarization and CaV1.3 channel opening change as a function of distance from this site is unclear. To explore this issue, NEURON simulations were performed using an SPN model.^66,68^To evoke a dendritic spike, 18 neighboring glutamatergic synapses were activated at 1 ms intervals in a 15-µm stretch of distal dendrite (Figure 6A). This synaptic stimulation produced a dendritic spike that pushed the local membrane potential to ∼-20 mV for about 20 msec; as expected, the depolarization of the somatic membrane potential was considerably smaller (Figure 6B). To provide more detail than the heat map of dendritic depolarization (Figure 6A), a plot of peak membrane potential as a function of distance from the soma was generated; it shows the strong attenuation with distance from the dendritic spike activation site (Figure 6C).

**Figure 6.**
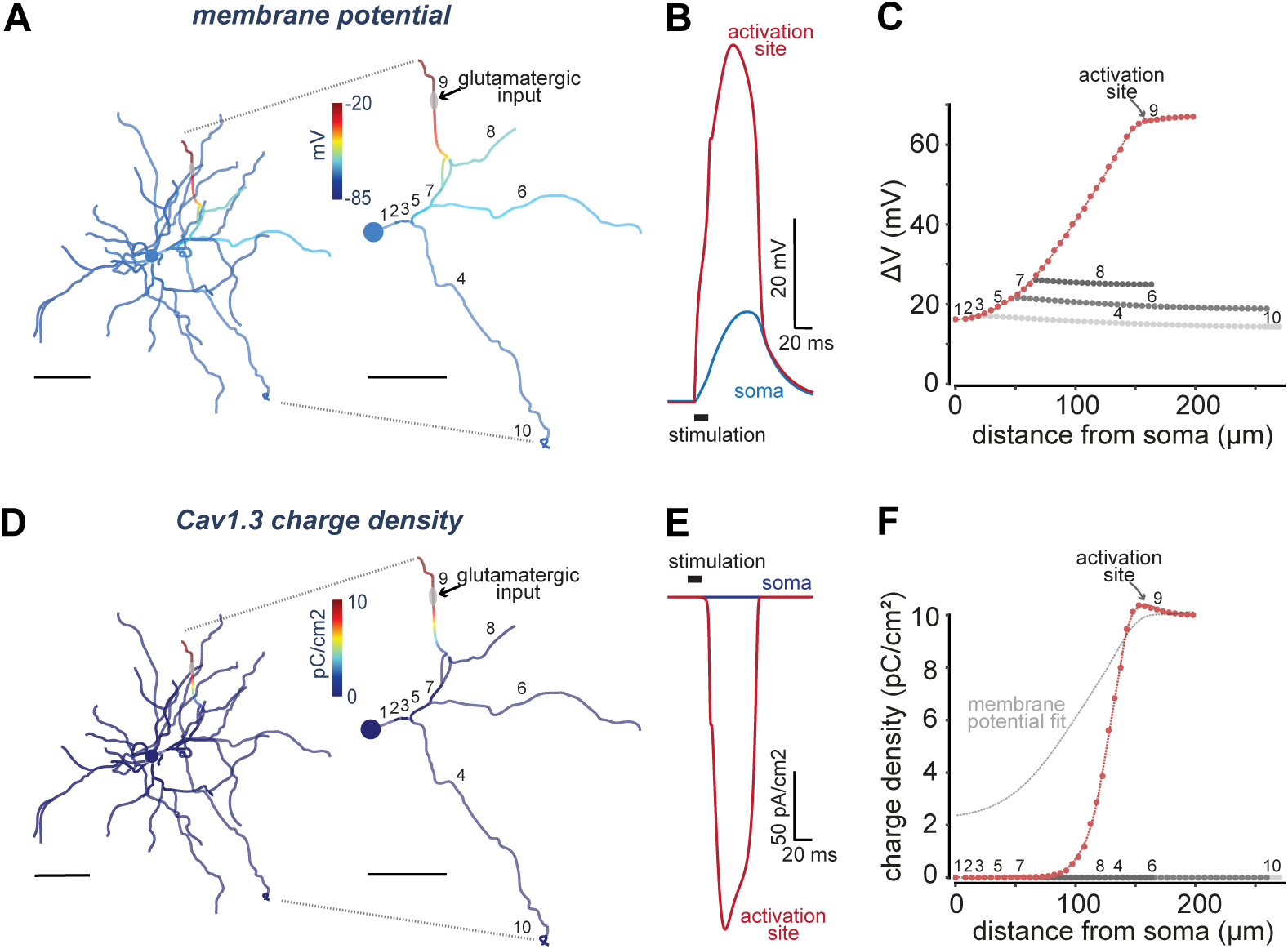
Computational modeling shows dendritic compartmentalization of membrane depolarization and CaV1.3 activation in response to dendritic spike generation. (A) Two-dimensional heatmap of a morphologically reconstructed dSPN illustrates peak membrane potential within the entire dendritic arbor in response to a dendritic spike. A dendritic spike is simulated at a distal location, as indicated by the arrow, by suprathreshold glutamatergic input (i.e. stimulating 18 neighboring synapses sequentially at 1 ms intervals in the distal-to-proximal direction). The zoomed-in section shows peak membrane potential within the dendritic path from the activation site to soma and in any major dendritic branches arising from this path (dendritic segments labelled 1-9). Scale bars here and in panel (D) are 50 μm. (B) Simulated synaptic potentials generated by clustered input at the site of activation and at the soma. (C) Peak change in membrane potential plotted as a function of path distance to the center of the soma for dendritic tree illustrated in the zoomed-in section of (A). (D) Equivalent 2-dimensional heatmap and zoomed-in section show total charge density (pC/cm^2^) due to activation of CaV1.3. (E) Local current density (pA/cm^2^) of CaV1.3 at the dendritic site of activation and soma. Charge density was calculated as the area under the current density plots over time. (F) Peak change in charge density plotted as a function of path length to the center of the soma at same locations as in (C). The fitted line showing membrane potential change along the main path from (C) has been superimposed to demonstrate the additional tuning effect the voltage gating properties of CaV1.3 has on the spatial profile of charge density within the dendritic tree. Data in (C) and (F) were fitted using univariate smoothing splines.

However, in agreement with previous work,^66,69^ distal to the site of spike generation, the peak membrane potential did not fall off significantly with distance (Figure 6C). To link this voltage measurement to the opening of CaV1.3 Ca^2+^ channels (and PDE1 activation), both a dendritic heat map and a detailed distance plot of CaV1.3 channel conductance was constructed (Figures 6D-6F). As expected from the non-linear relationship between membrane voltage and channel open probability, the CaV1.3 channel conductance fell off more rapidly in the proximal dendrite than did voltage; moreover, distal to the site of stimulation, CaV1.3 channels were strongly activated (Figures 6D-6F).

These results suggest that dendritic spikes will engage PDE1 to block cGMP signaling in dendritic regions that are located distally to the input site, but not dendritic regions that are more proximal to it, compartmentalizing NO-cGMP signaling in individual dendritic branches.

### Striatal NO signaling was disrupted in a PD model

Conventional forms of synaptic plasticity are lost in striatal SPNs following DA depletion with 6-hydroxydopamine (6-OHDA).^17,70–72^ It is not clear whether this deficit in PD models extends to NO-LTD. Indeed, PDE1 inhibition failed to induce synaptic depression in SPNs following an ipsilateral 6-OHDA lesion (Figures 7A and 7D). However, SNAP-induced LTD was normal (Figures 7B and 7D).

**Figure 7.**
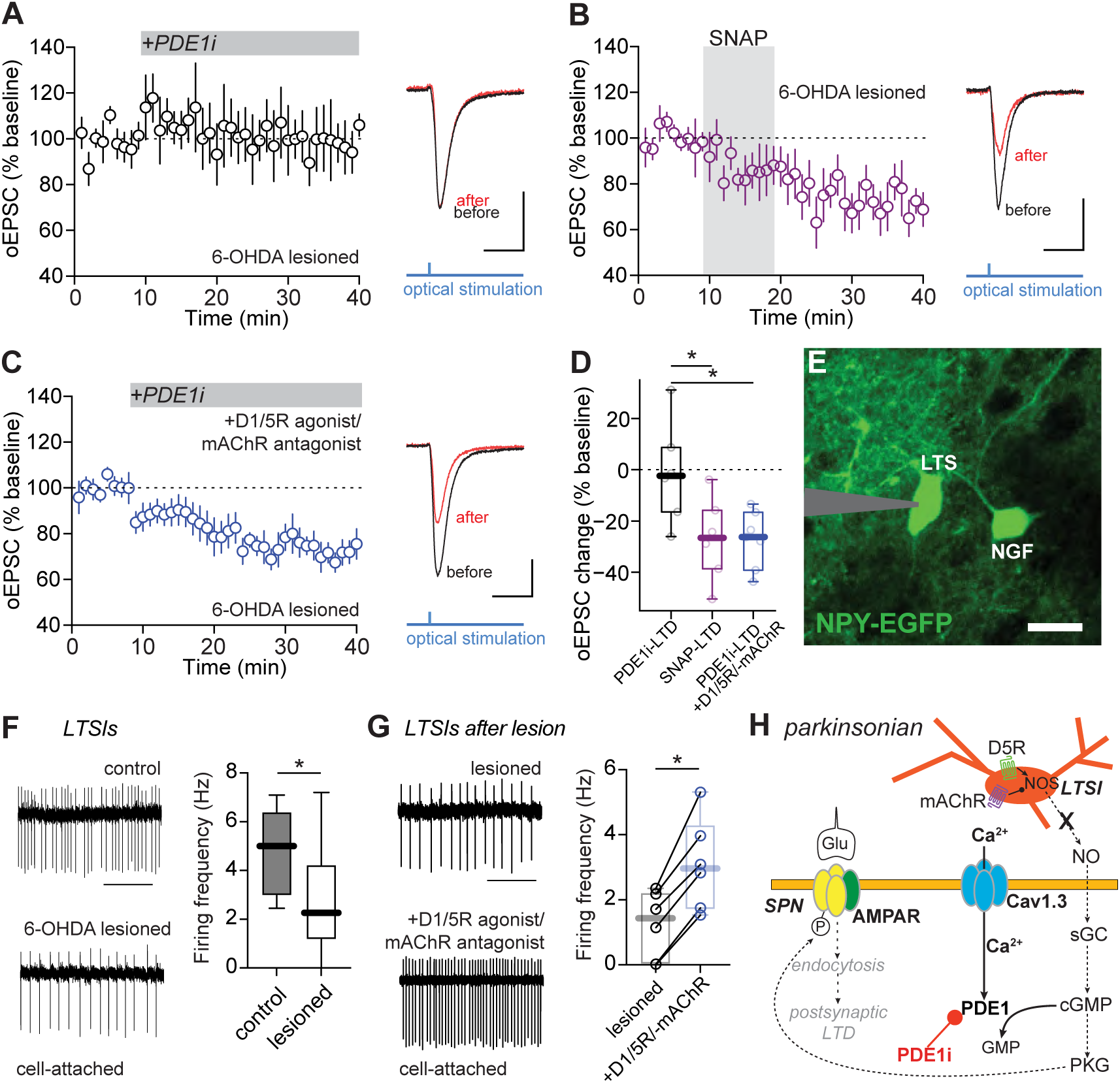
Endogenous NO release by LTSIs is impaired in parkinsonian mice and rescued by restoring the DA-ACh balance. (A) PDE1 inhibition fails to induce synaptic depression at corticostriatal synapses in iSPNs of 6-OHDA lesioned mice (n = 6 neurons from 3 mice). In panels (A-C), time course data represented as mean ± SEM are shown on the left. Sample EPSC traces before PDE1i application and from the last 5 min of recording are shown on the right. Scale bars are 100 pA x 20 ms. (B) SNAP-LTD is intact in iSPNs of 6-OHDA lesioned mice (n = 6 neurons from 4 mice). (C) In the presence of D1R/D5R-type agonist SKF 81297 (3 μM) and mAChR antagonist scopolamine (10 μM), PDE1i-LTD is restored (n = 7 neurons from 4 mice). (D) Box plot summary of data in (A-C) showing changes in oEPSC from the last 5 min of recordings. *p < 0.05, Mann-Whitney test. (E) Two-photon image of a low-threshold spiking interneuron (LTSI) in slices from NPY-EGFP mice recorded in cell-attached mode. Scale bar, 20 μm. (F) Left, examples of cell-attached recordings from LTSIs in slices from control or 6-OHDA lesioned NPY-EGFP mice. Right, box plot summary of firing rates of LTSIs (control, n = 7 neurons from 3 mice; 6-OHDA lesioned, n = 18 neurons from 4 mice). *p < 0.05, Mann-Whitney test. (G) Left, examples of cell-attached recordings from LTSIs from 6-OHDA lesioned NPY-EGFP mice before and after application of SKF 81297 and scopolamine (n = 6 neurons from 3 mice). Right, box plot summary of LTSI firing frequency in lesioned mice before and after application of D1-type agonist and mAChR antagonist. *p < 0.05, Wilcoxon test. (H) Schematic showing that in parkinsonian mice, lack of D5R signaling and enhanced mAChR signaling lead to reduction in endogenous NO release and postsynaptic LTD.

One possible explanation for the loss of NO-LTD following a 6-OHDA lesion is that DA depletion disrupted LTSI NO generation. The activity of LTSIs, and presumably their ability to generate NO, is modulated by both activation of D1-type DA receptors and muscarinic ACh receptors (mAChRs).^51,73,74^ As DA depletion results in elevated striatal ACh signaling,^49,75,76^ the ‘imbalance’ between these modulators could compromise the ability of LTSIs to generate NO. To test this hypothesis, the impact of PDE1 inhibition on synaptic transmission in iSPNs from 6-OHDA-lesioned mice was re-examined after ‘re-balancing’ neuromodulator tone by bath application of the D1-type agonist SKF 81297 (10 μM) and mAChR antagonist scopolamine (10 μM). These experiments focused on iSPNs because the robust D1R expression in dSPNs and their involvement in LTP induction may interfere with our examination of LTD.^17^ Re-balancing DA and ACh signaling restored the ability of PDE1 inhibition to induce LTD in iSPN from 6-OHDA lesioned mice (Figures 7C and 7D; p = 0.0141).

To better understand the effects of modulator re-balancing, patch clamp recordings were made from LTSIs identified by fluorescence in *ex vivo* brain slices from NPY-EGFP mice.^77^ LTSIs were readily distinguished from neurogliaform interneurons, the other type of interneurons that are labeled in NPY-EGFP mice, by their spontaneous activity and somatodendritic morphology (Figure 7E).^51,74,78^ In brain slices from 6-OHDA lesioned mice, the LTSI firing rate measured in cell-attached recordings was significantly lower than in controls (p = 0.0282) (Figure 7F). Bath application of the D1-type agonist SKF 81297 (10 μM) and mAChR antagonist scopolamine (10 μM), as above, normalized the spiking rate of LTSIs from lesioned mice (p = 0.0312) (Figure 7G). Taken together, these data suggest that the NO signaling deficit in PD models stems from the inhibition of LTSI spiking and a reduction in the intracellular Ca^2+^ signaling necessary for neuronal nitric oxide synthase (nNOS) activation (Figure 7H).^79^

## Discussion

The studies described here revealed that Ca^2+^ influx through depolarization-activated, CaV1.3 L-type Ca^2+^ channels, activates PDE1B, which potently suppresses constitutive NO/cGMP signaling and the induction of postsynaptic LTD at corticostriatal and thalamostriatal glutamatergic synapses on both iSPNs and dSPNs. As a result of this coupling, NO-LTD at glutamatergic synapses is state-dependent; that is, it is preferentially induced in dendritic regions that are relatively quiescent, contrasting it with other forms of activity-dependent synaptic plasticity in SPNs. The CaV1.3 channel dependence of PDE1B also helps to ensure that cGMP signaling and postsynaptic LTD are compartmentalized and do not compete with other forms of plasticity at glutamatergic synapses. Our studies also showed that a gain-of-function mutation in CaV1.3 channels linked to ASD attenuates NO-LTD induction in SPNs. Lastly, our work revealed that in an animal model of PD, constitutive NO signaling from LTSIs is disrupted, potentially contributing to a pathological distortion of the functional connectivity of SPNs and motor disability.

### Constitutively engaged NO-LTD is prominent in both iSPNs and dSPNs

Previously it was shown that NO released by LTSIs stimulates GC and cGMP signaling in SPNs, inducing LTD at glutamatergic synapses.^18,23,43^ In contrast to presynaptically expressed eCB-LTD, NO-LTD is expressed postsynaptically and mediated by endocytosis of AMPARs (shown here and also in ref. ^18^). Inhibition of PDE1B induced LTD in both iSPNs and dSPNs. Furthermore, unlike eCB-LTD at SPN glutamatergic synapses, NO-LTD can occur at both corticostriatal and thalamostriatal axospinous synapses. It also was expressed at axoshaft synapses formed by PFN terminals. Thus, essentially all glutamatergic synapses on DLS SPNs are subject to NO-LTD and regulation by PDE1B.

Precisely how cGMP signaling induces AMPAR endocytosis is unclear. Trafficking of AMPARs is regulated by a broad array of post-translational modifications (PTMs), including phosphorylation, nitrosylation, and interactions with an auxiliary subunit.^80^ Previously, NO-LTD in SPNs was postulated to be mediated by PKG, as it was blocked by the PKG inhibitor Rp-8-Br-PET-cGMPS.^18^ Indeed, at cerebellar parallel fiber-Purkinje cell synapses, PKG plays a key role in postsynaptic LTD induction by activating the protein phosphatase inhibitor G-substrate, which helps maintain PKC phosphorylation of GluA2 S880 necessary for retention of surface AMPA receptors (reviewed by ref. ^81^). Another potential target of cGMP signaling in SPNs is PDE2.^37^ PDE2 is stimulated by cGMP, but preferentially degrades cAMP.^82^ However, at a concentration selective for PDE2 (40 nM),^48^ Bay 60-7550 did not affect NO/cGMP-LTD in SPNs, suggesting that it was not mediated by PDE2 and suppression of PKA-dependent AMPAR trafficking into the synapse.^80^ NO-LTD in SPNs also differs from NMDAR-induced LTD,^80^ as insertion of Ca^2+^ permeable AMPARs was not a necessary step in its induction (it was unaffected by pre-incubation with NASPM). Additional work will be necessary to precisely define the mechanism underlying NO-stimulated AMPAR trafficking in SPNs.

Another important inference to be drawn from our studies was that cGMP signaling in SPNs was constitutive, as inhibition of PDE1 induced LTD in the absence of any other extrinsic stimulus. Autonomously active LTSIs are the principal source of striatal NO.^78^ Their activity elevates intracellular Ca^2+^ concentration – a necessary condition for activation of nNOS and NO generation.^83^ As NO readily crosses cell membranes, a single LTSI should be capable of influencing a region containing millions of synapses.^84^ Although there are no reliable methods for directly monitoring the spatiotemporal profile of NO produced by a single LTSI, the large axonal fields of LTSIs tile the dorsal striatum,^78^ suggesting that essentially all SPNs are exposed to a sustained, diffuse stimulation by NO. Moreover, GC, the nominal target of NO signaling, is robustly expressed and distributed throughout the dendritic tree of both major types of SPN.^38^

Using non-specific PDE inhibitors, a recent paper reported that cGMP signaling in cortical or thalamic terminals increased presynaptic Ca^2+^ influx and glutamate release.^85^ In contrast to the striatum, as shown by our results and others,^33,86^ the cortex has very low levels of PDE1B expression but robustly expresses PDE1A. It is possible that the reported cGMP effect on presynaptic release is regulated by PDE1A. Nevertheless, in our experiments, bath application of a membrane-permeable cGMP analog or an NO donor (SNAP) produced only synaptic depression that was attributable to a postsynaptic mechanism (see also^18^), suggesting that the dominant effect of cGMP on striatal glutamatergic synapses is depression.

### PDE1B activity was controlled by Ca^2+^ entry CaV1.3 channels

Three observations argue that PDE1B activity in SPNs is preferentially stimulated by Ca^2+^ entry through by CaV1.3 Ca^2+^ channels. First, the ability of PDE1B inhibition to induce LTD was disrupted by inhibiting CaV1 channels with the dihydropyridine negative allosteric modulator isradipine. Second, deleting the gene coding for the pore-forming subunit of CaV1.3 channels (*Cacna1d*) in adult SPNs (using viral delivery of Cre recombinase to SPNs in which *Cacna1d* was floxed) blocked the ability of PDE1B inhibition to induce LTD in SPNs. Lastly, in mice harboring a gain-of-function mutation in *Cacna1d* associated with ASD,^58^ the ability of the NO donor SNAP to induce LTD at corticostriatal synapses was significantly reduced. In contrast, eCB-LTD, which is stimulated by Ca^2+^ flux through CaV1.3 channels,^14,15,20,59,60^ was intact in mutant SPNs.

The potential biological logic of utilizing CaV1.3 channels to regulate PDE1B activity is three-fold. First, CaV1.3 channels are scaffolded by Shank isoforms close to SPN glutamatergic synapses, putting them in close proximity to the site of cGMP-induced plasticity.^87^ Second, Ca^2+^ entry through these channels is a positive modulator of eCB generation and presynaptic LTD.^14,15,20,59,60^ It makes excellent sense to make this one biological signal a negative modulator of a competing, functionally distinct form of synaptic plasticity. A third, distinctive feature of CaV1.3 channels is that they open at relatively hyperpolarized, sub-threshold membrane potentials. This feature may contribute to the compartmentalization of cGMP signaling by PDE1B. There are two subcellular regions of SPN to consider in this regard: proximal (∼<90 µm from soma) and distal dendrites. In the distal dendrites, spatiotemporally convergent activity at glutamatergic synapses can generate NMDAR-dependent spikes. These regenerative dendritic events create the conditions necessary for the induction of LTP at co-active glutamatergic synapses, which may allow SPNs to form associations between sensory and motor events.^7,66,67^ In agreement with others,^66^ our simulations suggest that these spikes strongly depolarize more distal regions of the dendrite, but rapidly decrement in amplitude in more proximal regions. The voltage-dependence of CaV1.3 channels ensures that PDE1B will be activated – and NO-LTD blocked – in the active dendritic branches. The sub-threshold voltage dependence of CaV1.3 channels should also ensure that more modest dendritic depolarization is sufficient to activate PDE1 and keep cGMP signaling restricted to dendrites that are below about -60 mV. Indeed, intracellular uncaging of cGMP in dendrites produced only a very localized LTD. In principle, this activity-dependent compartmentalization could extend to individual spines.^88^ Proximal dendrites of SPNs do not support synaptically triggered spikes, but spikes generated in the axon initial segment (AIS) decrementally propagate into those regions where they can trigger eCB-LTD or LTP.^17,89^ As in the more distal dendrites, the voltage-dependence of CaV1.3 channels should ensure that constitutive cGMP signaling is blunted when spikes invade this dendritic zone.

Thus, the coupling of CaV1.3 Ca^2+^ channels to PDE1B in SPNs establishes a reciprocity between signaling mechanisms controlling different forms of plasticity at glutamatergic synapses. Thus, when corticostriatal circuits activate during movement or goal-directed learning, SPN dendrites may be parceled into active regions where input-specific, activity-dependent forms of synaptic plasticity (eCB-LTD or LTP) is taking place and quiescent zones where counter-balancing synaptic depression mediated by cGMP signaling is happening.^63,90–93^ This interplay may be important to keep an SPN’s aggregate synaptic strength constant during associative learning, as well as to weaken synapses that are associated with extroceptive and interoceptive signals that are no longer relevant to ongoing control of goal-directed action.

### Possible behavioral functions of striatal NO signaling

How NO signaling figures into the role of the striatum in goal-directed action and habit is unclear. One obvious hypothesis is that NO-LTD drives the attenuation of infrequently activated synapses and in so doing facilitates ‘forgetting’ of associations that no longer are of value. In *Drosophila*, NO facilitates the rapid updating of memories, possibly by attenuating outdated associations.^94^ Consistent with a role in reversal learning, LTSIs integrate information coming from cortical, thalamic, and midbrain structures involved in evaluating and updating behavioral strategies. For example, they are richly innervated by neurons in the orbitofrontal cortex (OFC)^95^ which are critical to successful reversal learning,^96–98^ and by neurons in the anterior cingulate cortex (ACC), which are important to adapting motor plans.^99,100^ LTSIs also are linked to thalamostriatal circuits. In particular, activation of ChIs by thalamic PFN neurons inhibits LTSIs through both disynaptic GABAergic and muscarinic signaling pathways.^74,101,102^ Moreover, LTSIs have reward-associated activity that decreases acquisition of a goal-directed task.^103^ Lastly, LTSI activity and NO synthesis are enhanced by striatal release of dopamine,^73,74,104^ which is critical for reward-based reversal learning.^105–107^

Another potential clue about the potential role of NO signaling in regulating behavior comes from the recognition that mutations in CaV1.3 channels are associated with ASD. ASD patients commonly perseverate and have difficulty with reversal learning.^108–110^, Whole-exome sequencing of ASD patients revealed several risk-conferring mutations in *Cacna1d*.^111^ All these mutations, including the G407R mutation studied here, give rise to a gain of function in heterologous expression systems.^58,112^ Although many regions of the brain are likely to be affected by these mutations, the striatum is high on this list.^113,114^ For example, corticostriatal connectivity is elevated in ASD children.^115^ In *Cacna1d*^G407R/+^ mice, there was clear evidence of enhanced dendritic Ca^2+^ entry when SPN dendrites were depolarized.

Moreover, the induction of NO-LTD at corticostriatal synapses was disrupted in *Cacna1d*^G407R/+^ mice. Thus, it is reasonable to hypothesize that ASD-linked mutations in *Cacna1d* gene cause excessive CaV1.3 channel opening, increased PDE1B activity and blunted NO-LTD, leading to a deficit in suppression of old, contextually inappropriate associations and impaired behavioral flexibility.

### Dysregulation of NO-cGMP signaling in PD

Disruption of striatal NO-cGMP signaling has long been implicated in network dysfunction accompanying PD.^10^ Our studies provide new insight into this connection. A key feature of PD is the loss of the striatal dopaminergic innervation.^116^At the same time, striatal cholinergic signaling is elevated in the PD state because ChIs, which are normally suppressed by DA release, are dis-inhibited.^76,117–120^ Both neuromodulators strongly influence LTSIs: DA normally increases their basal spiking rate and promotes NO release,^73,74,121^ whereas ChIs inhibit LTSIs.^74^As a consequence, the PD state should be accompanied by lower autonomous LTSI spiking rates and reduced NO production. Previous studies showing that 6-OHDA lesions of the dopaminergic system reduce striatal cGMP levels are consistent with this hypothesis.^122,123^ Indeed, following a similar lesion of the dopaminergic system, basal LTSI activity was significantly depressed and constitutive NO-mediated suppression of SPN glutamatergic synapses lost. Importantly, both the reduction in LTSI spiking and deficit in constitutive NO/cGMP signaling was reversed in brain slices from 6-OHDA lesioned mice by pharmacologically ‘re-balancing’ DA and ACh signaling. Thus, the deficits in behavioral flexibility and reversal learning seen in models of PD^105^ and in PD patients^124^ could stem in part from a suppression of striatal NO signaling. The observation that NO signaling and its regulation of synaptic plasticity was effectively alleviated by the combination of a D1R agonist and mAChR antagonist has clear-cut implications for PD treatment regimens.

## Acknowledgments

We thank the members of the Surmeier lab and Dr. Richard B. Silverman for their support and comments on the manuscript. We thank Dr. Paul T. Schumacker and the Northwestern Transgenic & Targeted Mutagenesis Core Facility for their help in generation of the CaV1.3 floxed mice. We also thank Lundbeck for generously providing the PDE1B inhibitor AF64196. This work is supported by the JPB foundation (to D.J.S.), NS34696 (to D.J.S.), William N. and Bernice E. Bumpus Foundation (to D.J.S and to S.Z.), DOD W81XWH-18-1-0777 (to A.C. and D.J.S.), NIH/NIMH R01MH099114 (to A.C. and D.J.S.), Simons Foundation (to A.C.), NIH/NIGMS R35 GM131788 (to R.B.S.), NIH/NIGMS RO1 GM053395 and NIH/NINDS R35 NS111600 (to G.C.R.E-D.).

## Author Contributions

S.Z., D.J.S., S.O., and A.C. designed the experiments. S.Z. performed all electrophysiological and imaging experiments (with S.O. and D.W.’s help) and data analysis. S.O., J.X. and A.C. generated the *Cacna1d*^G407R/+^ mice. V.R.J.C conducted the computational modeling. T.T. and Z.X. performed the biochemical experiments and data analysis. A.T. helped with viral injection. H.T.D., C.T.R. and R.B.S. generated the nNOS inhibitor. H.K.A. and G.C.R.E-D. generated DEAC450-cGMP. S.Z. and D.J.S. prepared the manuscript. All authors approved the final version.

## Experimental Procedures

### Animals

All procedures were approved by the Northwestern Institutional Animal Care and Use committee. Mice (6-14 weeks of age) used in *ex vivo* experiments (except Figures 2E, 3H-3J and 7E-7G) were C57Bl/6 hemizygous mice expressing tdTomato or eGFP under control of Drd1a or Drd2 receptor regulator elements (RRID: MMRRC_030512-UNC and RRID: MMRRC_000230-UNC, backcrossed to C57BL/6 background). In some experiments, these mice were crossed with Thy1-ChR2-YFP mice (B6.Cg-Tg(Thy1-COP4/EYFP)18Gfng/J, RRID:IMSR_JAX:007612), *Cacna1d*^G407R/+^ mice (generated in-house), or Tg(Grp-cre)KH288Gsat (MMRRC, RRID: MMRRC_031183_UCD). In Figure 2E, ChAT-Cre x Ai14 mice (cross of RRID: IMSR_JAX:006410 and RRID: IMSR_JAX:007908) were used to identify ChIs. In Figures 3H-3J, CaV1.3^fl/fl^ mice generated in-house were used. In Figures 7E-7G, NPY-EGFP mice (RRID: IMSR_JAX:006417) were used to identify LTSIs. Both male and female mice were used for experiments.

### Brain slice preparation

The mice were anesthetized with intraperitoneal injection of a mixture of ketamine (100 mg kg^-1^) and xylazine (7 mg kg^-1^) and perfused transcardially with ice-cold cutting solution containing (in mM): 110 choline chloride, 25 NaHCO3, 1.25 NaH2PO4, 2.5 KCl, 0.5 CaCl2, 7 MgCl2, 11.6 sodium ascorbate, 3.1 sodium pyruvate and 5 glucose (305 mOsm L^-1^). Parasagittal brain slices (280-μm thick) were cut by a Leica vibratome (VT1200S, Leica Biosystems, Wetzlar, Germany) and then incubated for 40 min at 34°C in artificial cerebrospinal fluid (ACSF) containing (in mM): 124 NaCl, 3 KCl, 1 NaH2PO4, 2.0 CaCl2, 1.0 MgCl2, 26 NaHCO3 and 13.89 glucose, after which they were stored at room temperature for at least 30 min before recording. All external solutions were oxygenated with carbogen (95%CO2/5%O2) the entire time.

### Electrophysiology

Individual slices were transferred to a recording chamber and continuously superfused with ACSF (2-3 ml/min, 31-32°C). D1-Tdtomato- or D2-eGFP-expressing SPNs in the striatum were first identified with an Olympus BX-51-based two-photon laser scanning microscope (Ultima, Bruker Nano fluorescence). Whole-cell patch clamp was then performed in identified SPNs, aided by visualization with a 60X/0.9NA water-dipping objective lens and a ½” CCD video camera (Hitachi) imaged through a Dodt contrast tube and a 2x magnification changer (Bruker). All EPSC recordings were acquired in the presence of picrotoxin (50 μM) in the bath to suppress GABAA-mediated currents. For cGMP-mediated LTD recordings, patch pipettes (3-4 MΩ resistance) were loaded with an internal solution containing (in mM): 120 CsCH3SO3, 5 NaCl, 0.25 EGTA, 10 HEPES, 4 Mg-ATP, 0.3 Na-GTP, 10 TEA, 5 QX-314 (pH 7.25, osmolarity 280-290 mOsm L^-1^). SPNs were held at -70 mV (except in Figures 3F and 3G). For eCB-LTD experiments, patch pipettes (3-4 MΩ resistance) were loaded with an internal solution containing (in mM): 126 CsCH3SO3, 8 NaCl, 10 HEPES, 2.9 QX-314, 8 Na-phosphocreatine, 0.3 Na-GTP, 4 Mg-ATP, 0.1 CaCl2, 1 EGTA (pH 7.2-7.3, osmolarity 285-290 mOsm L^-1^).^19^ SPNs were held at -50 mV (Figure S8A) or -60 mV (Figures 4I, 4J and S8B) throughout the recordings. For cell-attached recordings of ChIs and LTSIs, interneurons were identified by their fluorescence, distinctive somatodendritic morphology and presence of spontaneous firing. Patch pipettes (∼4 MΩ resistance) were loaded with a solution containing (mM): 115 K-gluconate, 20 KCl, 1.5 MgCl2, 5 HEPES, 0.2 EGTA, 2 Mg-ATP, 0.5 Na-GTP, 10 Na-phosphocreatine (pH 7.25, osmolarity 280-290 mOsm L^-1^). All the recordings were made using a MultiClamp 700B amplifier (Axon Instrument, USA), and signals were filtered at 2 kHz and digitized at 10 kHz. Voltage clamp protocols and data acquisition were performed by *Prairie View* 5.3 (Bruker). Data were included if the series resistance (<20 MΩ) changed less than 20% and the holding current changed less than 100 pA over the course of the experiment. EPSCs were evoked every 20 s by single 0.3-ms pulses of whole-field LED illumination (oEPSC) or by paired electrical pulses (50 ms inter-stimulus interval) from a concentric bipolar electrode placed in Layer 5 of the overlying cortex. LED intensity and stimulus current were adjusted to yield EPSC amplitudes between 100-400 pA. EPSC amplitudes were normalized to the average EPSC of the baseline recording (first 5 or 8 minutes). The three normalized EPSCs from every minute were averaged to provide each data point in the LTD time course graphs.

### Whole-cell cGMP uncaging

The internal solution was supplemented with 75 μM DEAC450-cGMP. To avoid unintended photolysis of DEAC450-cGMP, internal solution preparation and patching were performed under red-light (630 nm) or orange-light (550 nm) illumination within a dark room. At the time of uncaging, patched cells have been filled with DEAC450-cGMP for at least 15 min. Whole-cell intracellular uncaging of DEAC450-cGMP was achieved with 200-ms pulses of 470/30 nm LED light (pE-100, CoolLED) at an intensity of 11.3 mW/mm^2^ measured at the objective lens (with a field of view of 366 μm diameter). For LTD induction, a total of five LED uncaging light pulses were applied at 1-min interval.

### Dual-color uncaging of glutamate and cGMP

The internal solution was supplemented with 75 μM DEAC450-cGMP and 50 μM Alexa Fluor-594 (Thermo Fisher Scientific). To avoid unintended photolysis of caged cGMP, the recorded SPN was visualized using a 1040-nm excitation laser (Chameleon-Ultra1, Coherent) from a two-photon laser scanning microscope (Ultima, Bruker) and a 60X/0.9 NA objective (Olympus). MNI-glutamate (5mM, Tocris), dissolved in HEPES-buffered ACSF, was superfused over the recorded region through a syringe pump (SP100i, World Precision Instruments), and a three-dimensionally localized volume of glutamate was photolysed by 1ms exposures from a separate two-photon-excitation laser (Verdi/Mira, Coherent) tuned to 720 nm as previously reported.^7^ Baseline 2pEPSC was recorded and averaged from four coplanar dendritic spines (∼50-60 μm from soma), each averaged 4 times (10 s inter-sweep interval). Localized dendritic uncaging of DEAC450-cGMP was initiated with 100-ms exposure of collimated 473 nm laser beam (Aurora, Prairie Technologies) at a light intensity of <28 mW/mm^2^. A series of five 473 nm laser spots (each spot being ∼7 μm in diameter, 100 ms in duration) was applied to cover a dendritic segment of ∼35 μm. Similar to the whole-cell cGMP uncaging, five uncaging epochs were applied at 1-min interval to induce LTD. At the end of each experiment, a maximum projection image (199.2 μm x 199.2 μm) of the SPN was acquired with 0.389 μm x 0.389 μm pixels, 1 μm z-steps, and 4 μs/pixel dwell time.

### Two-photon laser scanning Ca^2+^ imaging

The pipette solution (EGTA omitted) was supplemented with 100 μM calcium-sensitive dye Fluo-4 or Fura-2 (Thermo Fisher Scientific) and 50 μM calcium-insensitive dye Alexa Fluor 568 (Thermo Fisher Scientific). After whole-cell recording configuration was established, cells were allowed to equilibrate with dyes for at least 15 min before imaging. The recorded SPN was visualized using 810 nm (for Fluo-4 imaging) or 780 nm (for Fura-2 imaging) excitation (Chameleon-Ultra1: Coherent). Dendritic structure was visualized by the red signal of Alexa Fluor 568 detected by a Hamamatsu R3982 side-on photomultiplier tube (PMT, 580-620 nm). Calcium transients, as signals in the green channel, were detected by a Hamamatsu H7422P-40 GaAsP PMT (490-560 nm, Hamamatsu Photonics, Japan). Signals from both channels were background subtracted before analysis. Line scan signals were acquired with 128 pixels per line resolution and 10 μs/pixel dwell time along a dendritic segment. Ca^2+^ signals were quantified as the area of increase in green fluorescence from baseline normalized by the average red fluorescence (Λ1G/R) or the average baseline green fluorescence (Λ1F/F0).^22^

### Stereotaxic viral injection

Mice (8-12 weeks old) were anesthetized using an isoflurane precision vaporizer (at 5% isoflurane during induction and 2% isoflurane during maintenance phase) and positioned in a stereotaxic frame (David Kopf Instruments). Mice were administered with meloxicam (METACAM®, 0.1mg/kg, s.c., Covetrus) before surgery. After the skin and fascia were retracted to reveal the skull, a small hole was drilled, and an injection needle was slowly inserted. For studying synaptic responses at corticostriatal synapses, 0.15 μL AAV5-hSyn-hChR2(H134R)-EYFP (Addgene #26973) was injected into the M1 motor cortex at the following coordinates (mm relative to Bregma): ML 1.60, AP 1.15, DV 1.55. For studying synaptic responses at PFN-SPN synapses, 0.1 μL Cre-off virus AAV9-floxed-cre off-hEF1a-ChR2-EYFP-WRE-pA (Virovek, USA) was injected into the PFN of KH288 Grp-Cre mice at the following coordinates (mm relative to Bregma): ML 0.64, AP -2.23, DV 3.20. For RiboTag experiments, AAV9-EF1a-DIO-Rpl22l1-myc-FLag-2A-tdTomato (Virovek, USA) was injected into the striatum of A2a-Cre mice at the following coordinates (mm relative to Bregma): ML 2.3, AP1, DV3.2. The injection needle was left in place for 5 min to allow tissue absorption of the virus, and then withdrawn. The mice were then sutured up and left on a heating pad until recovery from anesthesia. Experiments were performed three weeks (for Cre-independent infection) or four to five weeks (for Cre-dependent infection) after viral injection.

### RiboTag profiling

The AAV for expressing RiboTag under a Cre-dependent promoter (AAV9-EF1a-DIO-Rpl22l1-Myc-DDK-2A-tdTomato-WPRE, titers 2.24 +10^13^ viral genome/mL) was stereotaxically injected into the striatum of A2a-Cre or A2a-Cre x CaV1.3^fl/fl^ mice as described above. Four weeks after injection, mice were sacrificed and the striatal tissue expressing RiboTag was dissected and frozen at -80°C. RiboTag immunoprecipitation was carried out as previously described ^49^. Briefly, tissue was homogenized in cold homogenization buffer (50 mM Tris at pH 7.4, 100 mM KCl, 10 mM MgCl2, 1 mM dithiothreitol, 100 μg/ml cycloheximide, protease inhibitors, and recombinant RNase inhibitors, 1% NP-40). Homogenates were centrifuged at 10,000x g for 10 minutes, the supernatant was collected and precleared with Protein G magnetic beads (Thermo Fisher Scientific) for 1 hour at 4°C, under constant rotation. Immunoprecipitations were carried out with anti-Flag magnetic beads (Sigma-Aldrich) at 4°C overnight with constant rotation, followed by four washes in high salt buffer (50 mM Tris at pH 7.4, 350 mM KCl, 10 mM MgCl2, 1% NP-40, 1 mM dithiothreitol, 100 μg/ml cycloheximide). RNA was extracted using RNA-easy Micro RNA extraction kit (QIAGEN) according to manufacturer’s instructions.

### Quantitative real-time PCR

RNA was extracted from the dissected striatum tissue using RNAeasy micro kit (QIAGEN). Complementary DNA (cDNA) was synthetized by using the Superscript IV VILO Master Mix (Applied Biosystems) and pre-amplified for 9 cycles using TaqMan PreAmp Master Mix and pool of TaqMan Gene Expression Assays (Applied Biosystems). The resulting product was diluted and then used for PCR with the corresponding TaqMan Gene Expression Assay and TaqMan Fast Advanced Master Mix. Data were normalized to hypoxanthine phosphoribosyltransferase 1 (HPRT) by the comparative the 2^−ΔΔCt^ method.^125^ The following TaqMan probes were used for PCR amplification of genes: HPRT, Mm03024075_m1; CaV1.2, Mm01188822_m1; CaV3.1, Mm00486549_m1; and CaV1.3, custom made (upper CCTGATTATTTTTACAGTGGAG, lower TCCATTCCTAACGTAAGCAT, probe ATAGCGTATGGACTGTTGCTG). Experimental Ct values were normalized to HPRT values using the formula: ΔCt = Ct (CaV1.2 or CaV1.3 or CaV3.1) – Ct (HPRT). Each ΔCt was then compared to the averaged ΔCt of the control group to obtain the ΔΔCt values: ΔΔCt = ΔCt-averaged ΔCt of control. The final expression levels were shown as 2^-ΔΔCt^.

### Immunostaining and confocal microscopy

Anesthetized mice were perfused transcardially with saline briefly (∼1 min) and then with ice-cold 4% paraformaldehyde (wt/vol) in 1x phosphate buffered saline (4% PFA-PBS). Brains were dissected out and post-fixed in 4% PFA-PBS overnight at 4°C. Immunostaining was performed using methods described previously.^126^ Briefly, sagittal or coronal slices (100 μm-thick) were cut using a Leica vibratome, permeabilized and blocked at 4°C in PBS including 5% normal goat serum and 0.2% Triton-X100 (NGS/PBST), incubated with rabbit monoclonal anti-PDE1B antibody (ab182565, Abcam) diluted 1:200 in NGS-PBST overnight at 4°C. After four washes in NGS/PBST, the slices were incubated with 1:1000 goat anti-rabbit Alexa Fluor 488 or 555 (A-11008 and A-21428, Invitrogen) for 2 hours at room temperature. Slices were washed with NGS-PBST four times and PBS once, mounted with VECTASHIELD Mounting Medium (Vector Laboratories), and viewed under an automated laser scanning confocal microscope (FV10i-DUC; Olympus). Images were adjusted for brightness, contrast, and pseudo-coloring in ImageJ (US National Institutes of Health).

### Western blotting

Striatum tissues from *Cacna1d*^G407R/+^ mice and wildtype siblings were collected and homogenized in N-Per Neuronal protein extraction buffer (Thermo Fisher Scientific) supplemented with Protease inhibitors (Halt Protease and Phosphatase Inhibitor Cocktail, Thermo Fisher Scientific). Samples were prepared in NuPAGE LDS Sample Buffer (Invitrogen) and separated by SDS-PAGE. Primary antibodies to PDE1B (1:1000, ab182565, Abcam) and GAPDH (1:1000, #2118, Cell Signaling,) were used for Western blotting. Target protein bands were visualized with horseradish peroxidase-conjugated secondary antibodies (Cell Signaling) and enhanced chemiluminescent reagent SuperSignal West Dura (Thermo Fisher Scientific). Intensities of the target protein bands were detected with Odyssey Fc (Li-Cor) and quantified in Image Studio (Li-Cor).

### Development of CaV1.3^fl/fl^ mice

The final targeting vector for conditional deletion of *Cacna1d* was purchased from EuMMCR (https://www.eummcr.org/search?q=PG00090_Y_4_H05&b=Search). The embryonic stem (ES) cell targeting and injection were performed by the Northwestern University Transgenic and Targeted Mutagenesis Laboratory. The targeting vector DNA was electroporated into PRX-B6N ES cells. Colonies that survived G418 selection were chosen and genotyped. After the correct targeting was confirmed, ES cells were injected into albino B6 (B6(Cg)-*Tyr*^c-2J^/J—000058 from The Jackson Laboratory) blastocysts and resulted in the generation of highly chimeric males (80-100% black in color). The chimera were then bred with albino B6 mice. The mutant offsprings were then mated with FLP recombinase transgenic mice to convert the unconditional allele into a conditional allele. The heterozygous conditional mice were then mated to generate homozygous (CaV1.3^fl/fl^) mice.

### Development of *Cacna1d*^G407R/+^ mice

The CRISPR/Cas9 gene editing system was used to generate the G407R amino acid substitution in the mouse *Cacna1d* gene.^127^ The glycine at position 407 (G407) is coded by two separate exons, with dinucleotide <u>GG</u> from exon 8 and a single nucleotide <u>A</u> from exon 9 making up the codon. A <u>GG</u> to <u>AG</u> mutation was introduced in exon 8 that results in a glycine to arginine substitution at this site (G407R) (Figure S7A). Single guide RNAs (gRNA) (Figure S7B), Cas9 mRNA and a single strand oligonucleotide donor (ss donor) with a single base mutation (G407 to R407) were microinjected into mouse zygotes. Two female founders with the positively identified G407R mutation were bred with wildtype C57/bl6J mice, and each of these founders have produced viable heterozygous and homozygous G407R mutant mice (see Figure S7C for a representative DNA sequencing chromatogram).

### Data acquisition, analysis and statistics

Electrophysiology and imaging data were acquired using PCI-NI6052E analog-to-digital converter card (National Instruments) and *Praire View* 5.3 (Bruker). Off-line analysis of electrophysiology data was performed using Clampfit 10.5 (Molecular Devices) and Origin 8 or 2020 (OriginLab). Time courses data were presented as mean ± SEM, and quantification data as non-parametric box-whisker plots. Off-line analysis of calcium imaging data was performed by custom-written Python script (available upon request), and statistical analysis performed by Prism 6 (GraphPad). The stated *n* indicates the number of cells (in electrophysiology experiments) or the number of dendrites (in calcium imaging experiments). Comparisons were made using Mann-Whitney test unless otherwise stated and differences with p < 0.05 are considered statistically significant.

### Modeling

The NEURON (Neuron 8.2; RRID:SCR_005393)^128^ + Python (Python Programming Language RRID:SCR_008394) model of a morphologically reconstructed SPN^68^ was built upon a previously established model.^66,129,130^ The model was essentially unchanged from the previously version (for details see Ref. ^68^; https://github.com/vernonclarke/SPNfinal/tree/v1.0; dx.doi.org/10.5281/zenodo.10162265). The number of compartments was increased slightly (∼820) and maximal values for the sigmoidally distributed CaV3.2 and CaV3.3 permeability reduced to 10^-6^ and 10^-7^ cm/s, respectively.

## RESOURCE AVAILABILITY

### Lead Contact

Further information and requests for resources and reagents should be addressed to the Lead Contact Dr. D. James Surmeier.

### Material Availability

Request for *Cacna1d*^G407R/+^ mice should be addressed to Dr. Anis Contractor; request for DEAC450-cGMP should be addressed to Dr. Graham Ellis-Davies; request for CaV1.3^fl/fl^ mice and all other materials should be addressed to Dr. D. James Surmeier.

### Data and Code Availability

The raw data are available with reasonable request to the Lead Contact Dr. D. James Surmeier. The custom-written Python scripts for analyzing imaging and physiology results are available at: https://github.com/ShenyuZhai/Data-analysis-codes. The NEURON + Python model of a reconstructed SPN is available at: https://github.com/vernonclarke/SPNfinal/tree/v1.0; dx.doi.org/10.5281/zenodo.10162265).

## Supplemental information

Supplemental information includes nine figures (Figures S1-S9).

**Figure S1.**
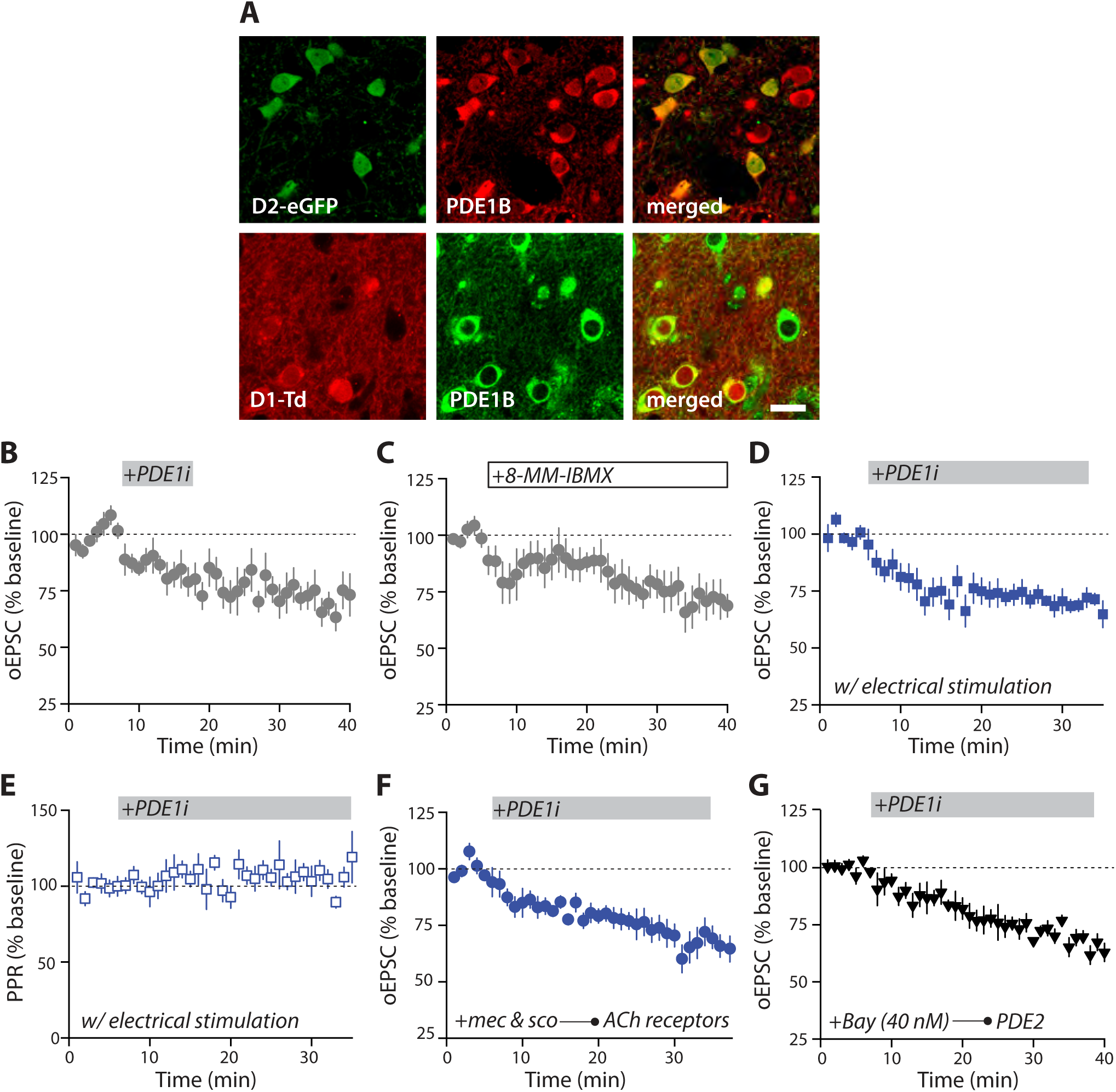
PDE1B regulates cGMP signaling in SPNs (related to Figure 1) (A) Representative confocal images showing colocalization of PDE1B immunoreactivity with D2-eGFP fluorescence (top) and with D1-tdTomato fluorescence (bottom). Scale bar is 20 μm. (B) A 10-min application of PDE1i induced persistent LTD at corticostriatal glutamatergic synapses (n = 7 neurons from 5 mice). In this panel and panels (C-G), normalized EPSC amplitudes are plotted over time and error bars indicate SEM. (C) A less selective PDE1 inhibitor 8MM-IBMX (10 μM), like AF64196, causes synaptic depression at corticostriatal glutamatergic synapses in SPNs (n = 8 neurons from 8 mice). (D) Synaptic depression could be induced by PDE1i when EPSCs were evoked by electrical stimulation with a concentric bipolar electrode placed at Layer 5 of overlaying cortex (n = 6 neurons from 6 mice). (E) Normalized paired-pulse ratio (PPR) of electrically evoked EPSCs was not significantly changed by PDE1i application. (F) Blocking cholinergic signaling with nicotinic antagonist mecamylamine (mec, 10 μM) and muscarinic antagonist scopolamine (sco, 10 μM) did not prevent PDE1i-LTD (n = 7 neurons from 4 mice). (G) Inhibition of PDE2 with Bay 60-7550 (Bay, 40 nM) did not prevent PDE1i-LTD (n = 6 neurons from 3 mice).

**Figure S2.**
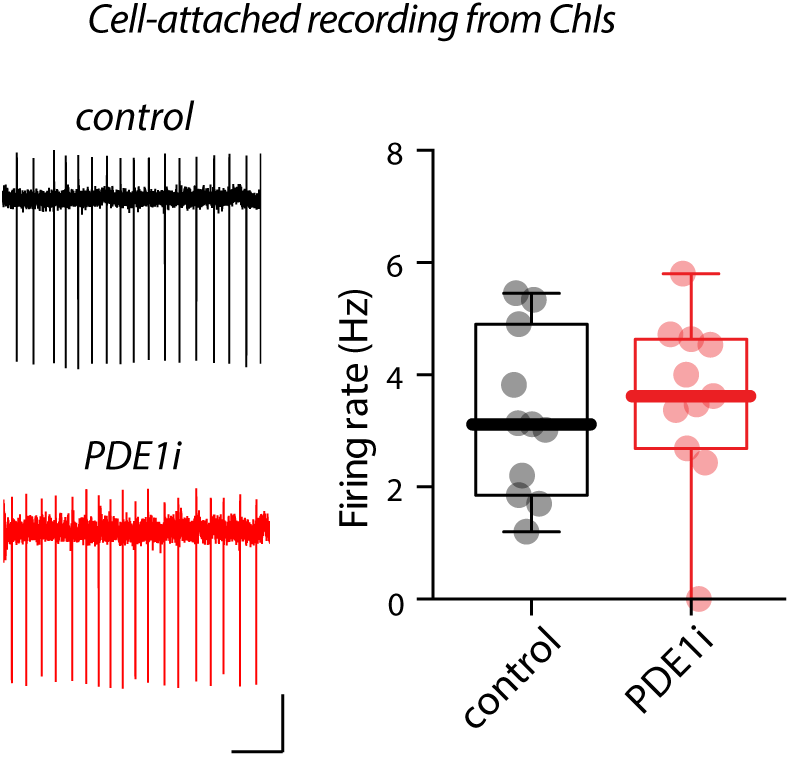
PDE1B inhibitor has no effect on ChI activity (related to Figure 2) Sample cell-attached recordings (left) and box plot summary (right) showing lack of effect of PDE1i on ChI firing rate. Scale bars represent 30 pA and 1 s. p > 0.05, Mann-Whitney test.

**Figure S3.**
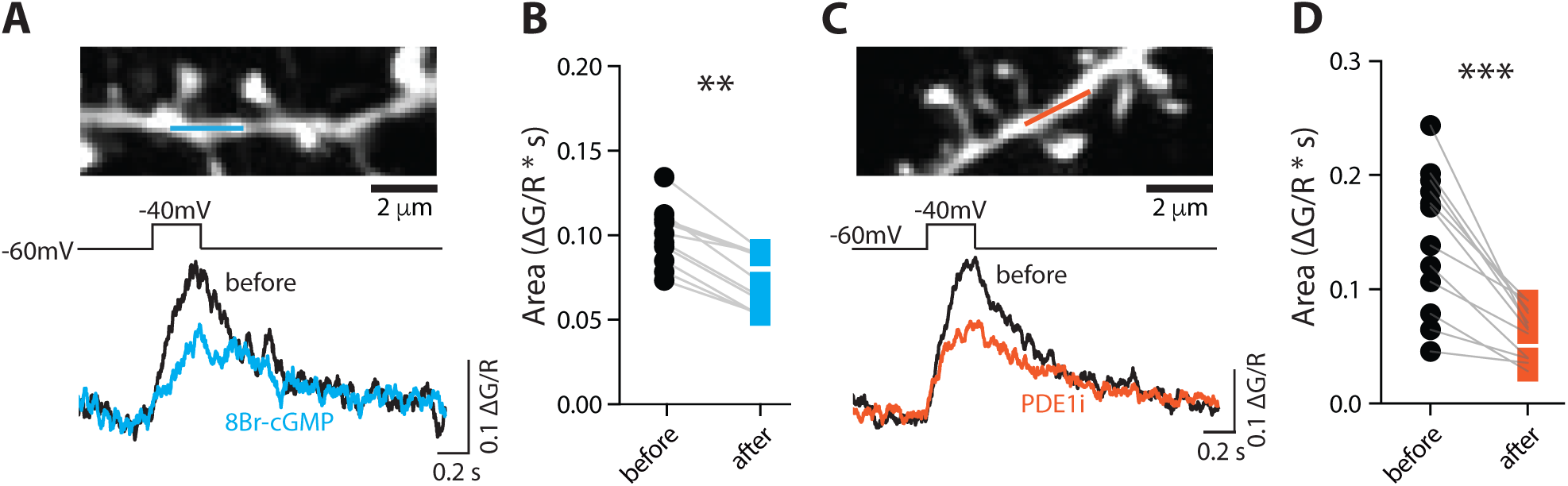
PDE1B inhibitor, as the cGMP analog, reduces dendritic calcium transient. (A) Top, two-photon laser scanning microscopy image of a dendritic segment from which a Ca^2+^ imaging line scan (indicated by the blue line) was performed. Middle, voltage step protocol used for optimally isolating L-type Ca^2+^ signals. Bottom, examples of fluorescence signals recorded before and after cGMP analog 8Br-cGMP (500 μM) application. Here, two-photon Ca^2+^ imaging was performed with Fluo-4 with excitation wavelength at 810 nm. (B) 8Br-cGMP reduced dendritic fluorescence transients evoked by voltage step protocol. n = 10 dendrites from 5 cells from 3 animals. (C and D) Inhibition of PDE1, like cGMP analog, negatively modulates L-type Ca^2+^ transients. Same as in (A) and (B) but with PDE1i. n = 12 dendrites from 6 cells from 3 animals. **p < 0.01, ***p < 0.001, Wilcoxon test.

**Figure S4.**
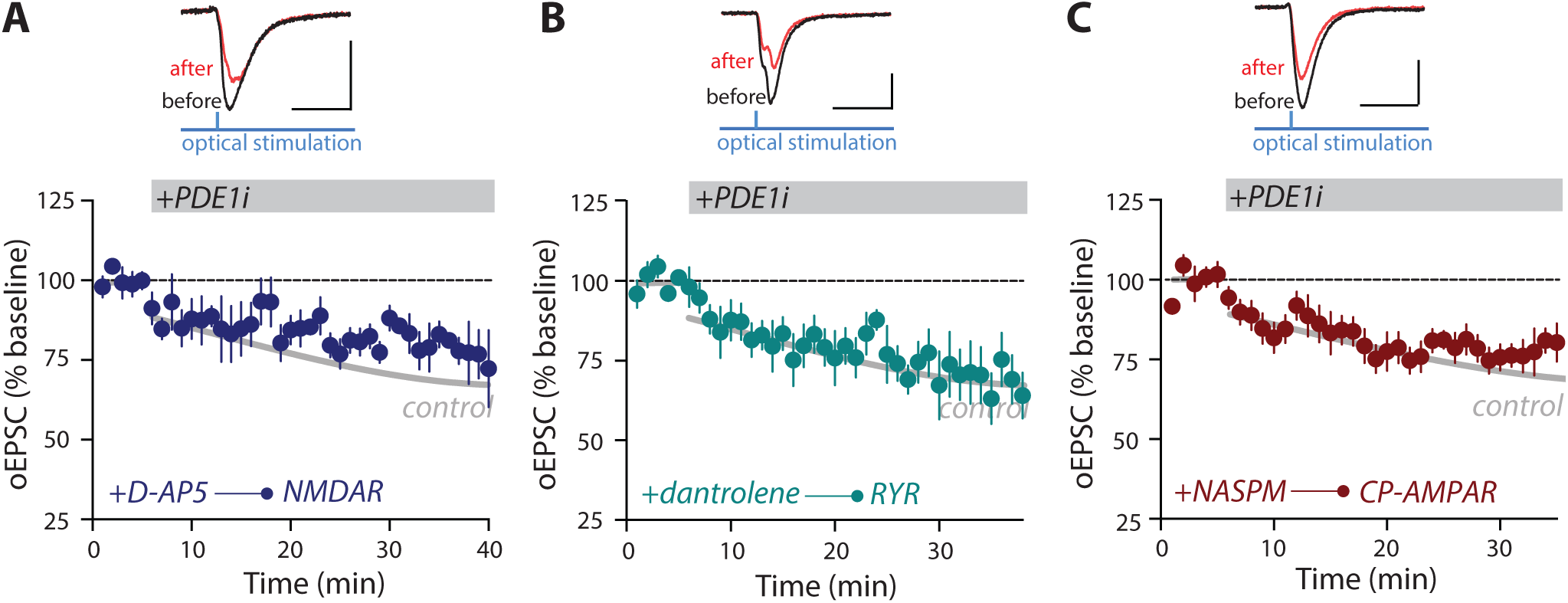
PDE1 activity does not depend on Ca^2+^ from NMDARs, ryanodine receptor-mediated Ca^2+^ release or Ca^2+^-permeable AMPARs (related to Figure 3) PDE1i-induced LTD in the presence of NMDAR antagonist D-AP5 (50 μM, n = 6 neurons from 4 mice) (B), ryanodine receptor antagonist dantrolene (10 μM, n = 6 neurons from 4 mice) (C), and Ca^2+^-permeable AMPAR antagonist NASPM (20 μM, n = 6 neurons from 4 mice) (D). In all panels, normalized EPSC amplitudes are plotted over time. Sample traces are shown at the top. The black ‘before’ traces show the average EPSC from 0-5 min and the red ‘after’ traces show the average EPSC from the last 5 min of recordings. Scale bars are 100 pA and 20 ms and error bars are SEM.

**Figure S5.**
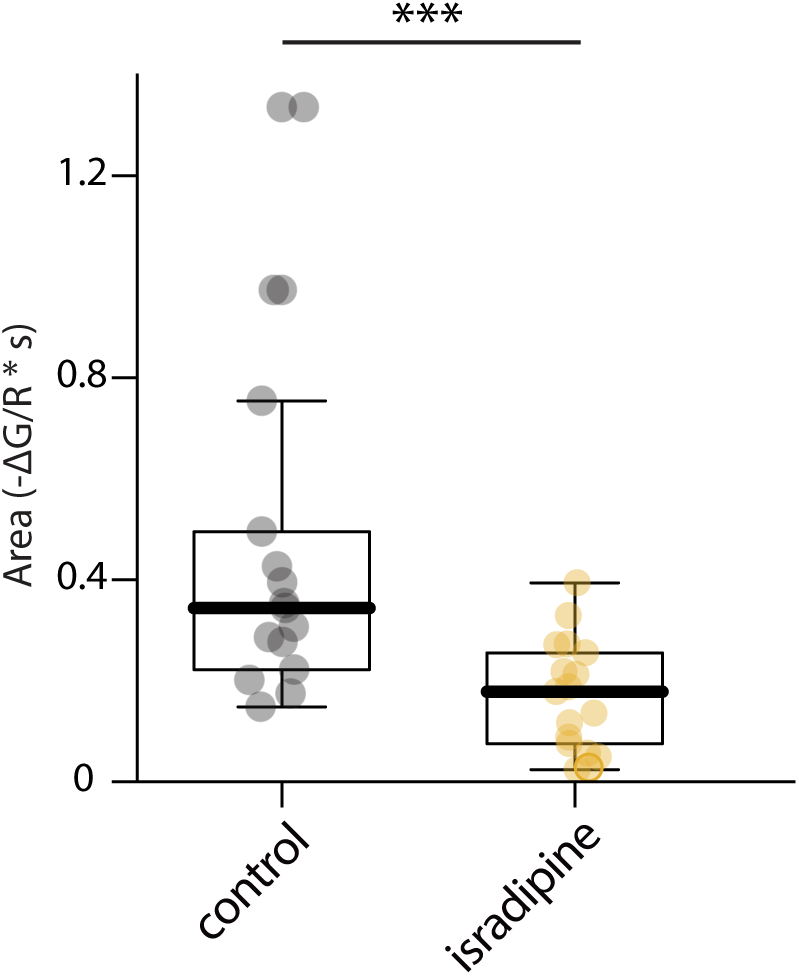
Dendritic Ca^2+^ transients evoked by depolarization to -50 mV are significantly blocked by isradipine, a negative allosteric modulator of L-type Cav1 Ca^2+^ channels (related to Figure 3). Box plot summary of dendritic Ca^2+^ signals evoked by depolarization of SPNs from -70 mV to -50 mV for 2.5 s in control condition or with bath application of isradipine (1 μM). Here, two-photon Ca^2+^ imaging was performed with Fura-2 with excitation wavelength at 780 nm. n = 17 dendrites from 5 cells that were from 3 animals. ***p = 0.0004, Mann-Whitney test.

**Figure S6.**
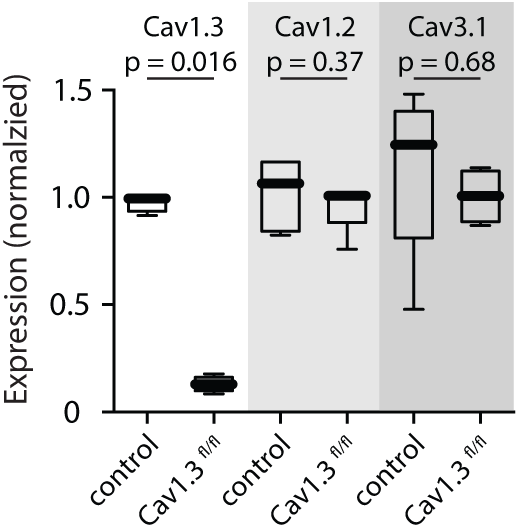
Validation of CaV1.3^fl/fl^ mice (related to Figure 3) RiboTag-based gene profiling confirms deletion of CaV1.3 in iSPNs of A2a-Cre x CaV1.3^fl/fl^ mice. Box plot showing expression levels of CaV1.3, CaV1.2 and CaV3.1 in A2a-Cre x CaV1.3^fl/fl^ mice or A2a-Cre control (see Experimental Procedures). n = 5 animals for A2a-Cre controls and n = 4 animals for A2a-Cre x CaV1.3^fl/fl^ mice. p values are indicated on the graph, Mann-Whitney test.

**Figure S7.**
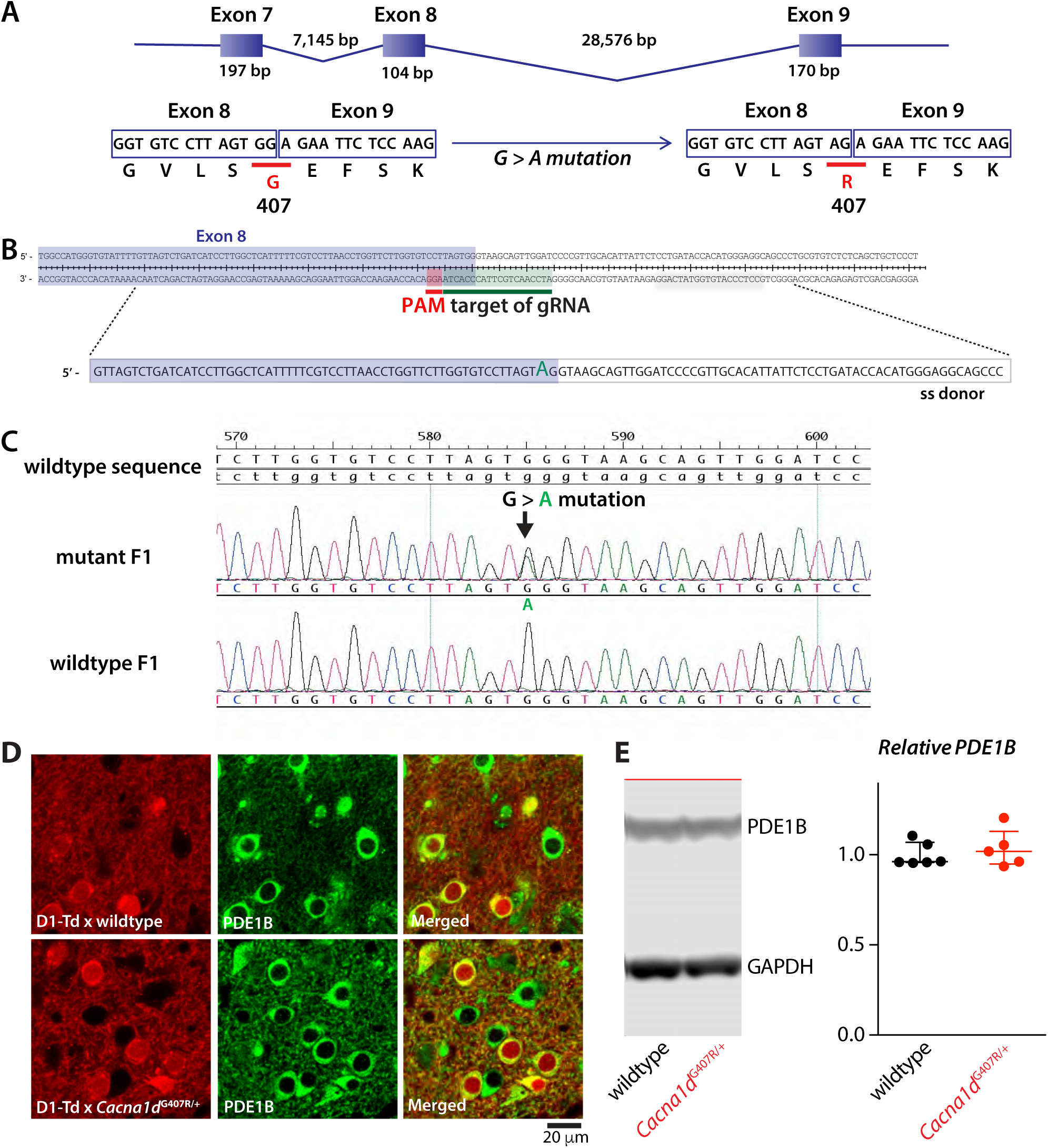
Generation and characterization of *Cacna1d*^G407R/+^ mice (related to Figure 4) (A) Schematic illustration of genomic structure of *Cacna1d* locus containing exons 7-9. Boxes and lines indicate exons and introns. G407 is coded by two exons: (GG) in exon 8 and (A) in exon 9. Introduction of a mutation in exon 8 to convert the dinucleotide <u>GG</u> to <u>AG</u> results in a glycine (G)-to-arginine (R) switch at position 407 (G407R). (B) Top, parts of exon 8 of the mouse *Cacana1d* gene and ensuing intron that contain the target of guide RNA (gRNA). The protospacer adjacent motif (PAM) sequence was highlighted in red. Bottom, the sequence of the single strand oligonucleotide (ss) donor is shown. (C) DNA sequencing chromatograms of a representative F1 generation mouse with a monoallelic G > A substitution (middle) and a wildtype littermate (bottom). The genomic DNA around the gRNA target site was PCR amplified and directly sequenced. (D and E) PDE1B expression in *Cacna1d*^G407R/+^ mice is indistinguishable from that in wildtype siblings. Confocal images (D) showed similar PDE1B immunoreactivity (green) in the dorsal striatum from wildtype or *Cacna1d*^G407R/+^ mice that express tdTomato under Drd1a receptor regulatory element (D1-tdT). Also shown by Western blotting (E), PDE1B protein levels were similar between *Cacna1d*^G407R/+^ mice and wildtype controls. Left, a representative Western blot showing PDE1B and GAPDH protein levels in wildtype and mutant. Right, summary plot of relative PDE1B levels normalized to GAPDH as the loading control (p = 0.7922, Mann-Whitney test; n = 6 wildtype and 5 *Cacna1d*^G407R/+^ mice).

**Figure S8.**
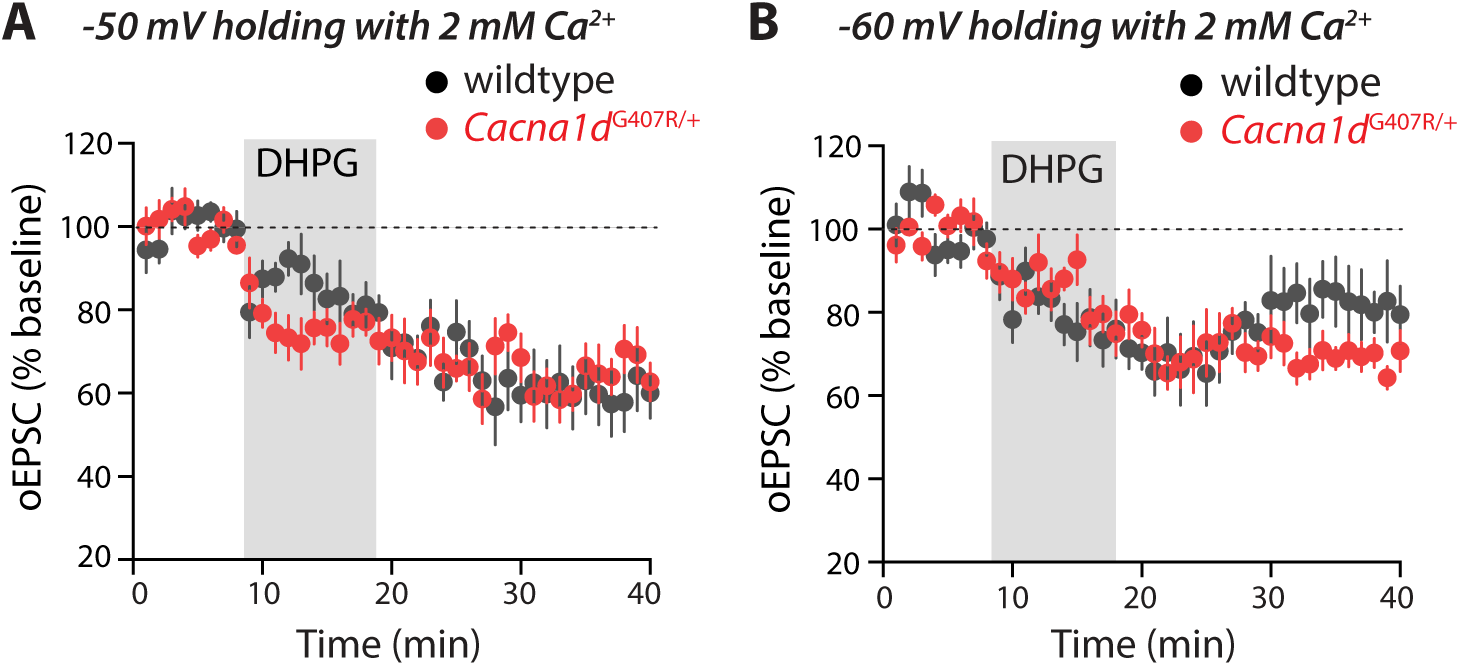
Endocannabinoid-induced LTD (eCB-LTD) in *Cacna1d*^G407R/+^ mice (related to Figure 4) DHPG-LTD in *Cacna1d*^G407R/+^ mice is indistinguishable from that in wildtype siblings. DHPG (50 μM) application for 10 min induced robust LTD in both *Cacna1d*^G407R/+^ mice and wildtype siblings when the postsynaptic dSPNs were voltage clamped at -50 mV (A) or -60 mV (B). In both panels, mean ± SEM are shown. In (A), n = 9 cells from 4 wildtype mice and n = 6 neurons from 3 *Cacna1d*^G407R/+^ mice; in (B), n = 7 neurons from 4 wildtype mice and n = 6 neurons from 4 *Cacna1d*^G407R/+^ mice.

**Figure S9.**
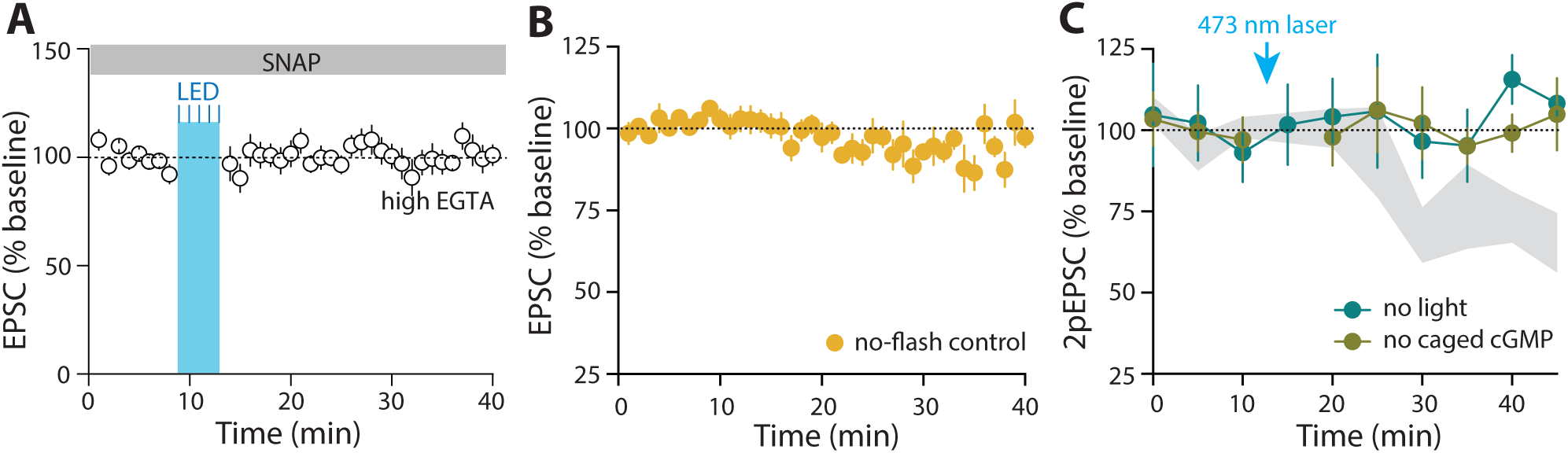
Control experiments with DEAC450-cGMP uncaging (related to Figure 5) (B) Pretreatment with SNAP (100 µM) occluded LTD induced by DEAC450-cGMP uncaging (n = 6 neurons from 4 animals). (C) Control experiments for cGMP uncaging with blue LED. Without blue LED flashes (‘no-flash control’), intracellular dialysis of caged cGMP did not alter EPSC amplitudes. Sample size is 6 neurons from 5 mice. (D) Control experiments for cGMP uncaging with 473 nm laser. Caged cGMP alone (‘no light’) or 473 nm laser alone (‘no caged cGMP’) had no effect on the amplitude of 2pEPSCs. The blue arrow indicates the timing of 473 nm laser illumination if there was one. The ‘cGMP+glu uncaging’ data from Figure 5F was shown as grey shade for comparison. No light, n = 4 neurons from 3 mice; no caged cGMP, n = 5 neurons from 3 mice.

